# The adiponectin analogue ALY688-SR attenuates diaphragm fibrosis, atrophy and mitochondrial stress in a mouse model of Duchenne muscular dystrophy

**DOI:** 10.1101/2023.05.22.541826

**Authors:** Catherine A. Bellissimo, Shivam Gandhi, Laura N. Castellani, Mayoorey Murugathasan, Luca J. Delfinis, Arshdeep Thuhan, Madison C. Garibotti, Yeji Seo, Irena A. Rebalka, Gary Sweeney, Thomas J. Hawke, Ali A. Abdul-Sater, Christopher G.R. Perry

**Affiliations:** School of Kinesiology & Health Science, Muscle Health Research Centre, York University, Toronto, ON M3J 1P3 Canada; Department of Biology, Muscle Health Research Centre, York University, Toronto, ON M3J 1P3 Canada; Department of Pathology and Molecular Medicine, McMaster University, Hamilton, ON, Canada

**Keywords:** Keywords, Duchenne muscular dystrophy, neuromuscular disease, dystrophin, weakness, atrophy, fibrosis, muscle, mitochondria, adiponectin, oxidative stress

## Abstract

Fibrosis is associated with respiratory and limb muscle atrophy in Duchenne muscular dystrophy (DMD). Current standard of care partially delays the progression of this myopathy but there remains an unmet need to develop additional therapies. Adiponectin receptor agonism has emerged as a possible therapeutic target to lower inflammation and improve metabolism in *mdx* mouse models of DMD but the degree to which fibrosis and atrophy are prevented remain unknown. Here, we demonstrate that the recently developed slow-release peptidomimetic adiponectin analogue ALY688-SR prevents fibrosis and fibre type-specific atrophy in diaphragm of D2.*mdx* mice treated from days 7-28 of age. ALY688-SR also lowered IL-6mRNA but increased IL-6 and TGF-β protein contents in diaphragm, suggesting dynamic inflammatory remodeling. ALY688-SR alleviated mitochondrial redox stress by decreasing complex I-stimulated H_2_O_2_ emission. Treatment also lowered *in vitro* diaphragm force production in diaphragm suggesting a complex relationship between adiponectin receptor activity, muscle remodeling and force generating properties during the very early stages of disease progression in D2.*mdx* mice. In tibialis anterior, the modest fibrosis at this young age was not altered by treatment, and atrophy was not apparent at this young age. These results demonstrate that short-term treatment of ALY688-SR partially prevents fibrosis and atrophy in the more disease-apparent diaphragm of young D2.*mdx* mice in relation to lower mitochondrial redox stress. These results provide a foundation for the exploration of slow-release adiponectin-based therapies to prevent fibrosis and atrophy in Duchenne muscular dystrophy.

## Introduction

Duchenne muscular dystrophy (DMD) is a neuromuscular disorder estimated to occur in approximately 1 in 5,000 live male births ^1^. Progressive muscle weakness in childhood usually leads to an early dependence on assistive devices as well as a reduced lifespan from respiratory or cardiac failure. Arising from an X-linked recessive mutation in dystrophin, a loss of cell membrane stability in striated muscles renders these cells susceptible to contraction-induced damage leading to repeated cycles of degeneration, systemic inflammation and fibrosis ^2–5^. While immunosuppression by glucocorticoids slows the progression of the disease, the considerable myopathy that remains combined with the deleterious side effects of this treatment underscores an unmet need for developing additional therapeutics.

Adiponectin is an adipokine that lowers inflammation ^6, 7^ and upregulates indices of fat oxidation in skeletal muscle ^8–10^. Transgenic overexpression of adiponectin ^11^ in C57BL/10.*mdx* mice improved whole-body measures of muscle function and reduced markers of inflammation and injury. Likewise, the orally administered small-molecule (non-peptide) adiponectin receptor (AdipoR) agonist, AdipoRon, reduced markers of inflammation and sarcolemmal damage in skeletal muscle from C57BL/10.*mdx* mice, stimulated genomic markers of regeneration and improved whole body assessments of locomotor function using treadmill, grip strength and wire tests ^12^. These effects were associated with general indices of reduced oxidative stress but the precise mechanism by which this occurred remains unknown. Also, the effects of AdipoR agonists on other hallmark signs of myopathy in *mdx* mice such as muscle-specific weakness, fibrosis, atrophy and the well-characterized state of metabolic stress (reviewed in ^13^) remain unknown.

Whilst adiponectin and AdipoRon are efficacious, the former is a large multimeric protein which requires extensive posttranslational modification making it expensive to synthesize and the latter lacks potency and specificity ^14, 15^. The critical receptor binding domain of adiponectin was identified and used to generate the peptidomimetic ADP355 which resulted in high specificity, was more potent than the native protein and more resistant to proteolytic degradation. ADP355, now known as ALY688, has been shown to activate adiponectin-specific signaling downstream of both AdipoR1 and AdipoR2 receptors ^15, 16^. ALY688 was also shown to attenuate inflammation in various inflammatory disorders such as dry eye and liver diseases, with reduced fibrosis also seen in the latter ^17, 18^. This anti-fibrotic effect is intriguing given the considerable fibrosis that occurs in DMD and murine models which may be mediated by inflammation and contribute to the progressive decline in muscle function ^19–21^. As metabolic dysfunction is also apparent in DMD and *mdx* mice (reviewed in ^13^), the recent finding that ALY688 improves metabolic function in skeletal muscle cells ^22^ and adipocytes ^23^ further supports consideration of this compound.

Importantly, for preclinical studies a slow-release formulation (ALY688-SR) was recently developed with improved pharmacokinetic properties enabling once daily injection (unpublished observation). The purpose of this study was to determine the effects of *in vivo* administration of ALY688-SR on fibrosis, atrophy, force production and mitochondrial bioenergetics in dystrophin-deficient muscle. Herein, we used D2.*mdx* mice – a model that demonstrates more robust muscle dysfunction compared to C57BL10.*mdx* mice ^24, 25^. We focused specifically on the early stages of disease progression by treating mice from 1-4 weeks of age given that symptoms first manifest in young children ^26^ and that we have previously characterized myopathy and mitochondrial stress in 4-week-old D2.*mdx* mice ^27, 28^. In the present study, we compared both diaphragm and limb muscles given their clinical relevance to respiratory and locomotor dysfunction in DMD. Here, we demonstrate that ALY688-SR attenuates fibrosis in both muscles and prevents atrophy in specific muscle fibre types from the diaphragm in relation to attenuated mitochondrial hydrogen peroxide (mH_2_O_2_) emission. Divergent responses in cytokine markers and reduced *in vitro* diaphragm force production were also observed. Collectively, these results suggest that adiponectin receptor agonism causes dynamic and complex muscle remodeling when administered during early stages of disease progression and at an age of rapid development. The longer-term effects of this early attenuation in fibrosis, atrophy and mitochondrial stress on force generation capacity warrants further investigation.

## Materials and Methods

### Animal care

Male D2.*mdx* mice were bred from an in-house colony established at York University (Toronto, Ontario) and treated from 7 to approximately 28 days of age. 3 week-old DBA/2J wildtype (WT) mice were ordered directly from Jackson Laboratories (Bar Harbor, MI, USA) due to low breeding performance experienced in-house as reported previously ^29^ and allowed to acclimatize for 7 days prior to sacrifice. Mice were housed with littermates until functional testing (commencing 48 hours prior to sacrifice) and then sacrifice (∼28 −30 days of age). Mice were maintained on a 12:12-h light-dark cycle while being provided access to standard chow and water ad libitum. Given small tissue size, multiple phases of breeding were required for sufficient masses for analyses. As a result, two phases of breeding and treatments were required. Mice were randomly assigned to the respective groups. All experiments and procedures were approved by the Animal Care Committee at York University (AUP Approval Number 2016-18) in accordance with the Canadian Council on Animal Care.

### ALY688-SR treatment

D2.*mdx* mice were treated with ALY688-SR (See ‘Acknowledgements’ for more information from Allysta Pharmaceuticals, Bellevue, WA, USA) at two different dosages; high dose (HD; 15mg/kg body weight/day) or low dose (LD; 3mg/kg body weight/day) dissolved in saline. A separate group of mice were treated with saline alone (VEH, vehicle). Treatment was provided via subscapular subcutaneous injections from 7 days of age to 28-30 days. Wildtype mice did not receive treatment. At 28-30 days of age, mice were anaesthetized with isoflurane vaporized in medical air (21% oxygen) at a 2L/min flow rate. *In situ* quadriceps force assessments were then performed in one limb followed by removal of the quadriceps from the other limb for mitochondrial assessments, diaphragm for in vitro force assessments and finally other tissues for further processing as described below. Muscle and visceral organs weights were normalized to tibia length. Given tissue limitations, several phases of breeding were performed. Tissue collected from the first phase were utilized in force, histological, mitochondrial measures (respiration, mH_2_O_2_ and calcium retention capacity) and western analyses (diaphragm and quadriceps only). A second phase was then collected and utilized for cytokine profiling, qPCR analyses and additional westerns (TA only). Serum was collected from both phase 1 and 2 and utilized for serum CK analysis. All animals were utilized in in vivo functional assays (grip strength, cage hang time and voluntary wheel running).

### RNA isolation & quantitative PCR

Samples were lysed using TRIzol reagent (Thermo Scientific, Waltham, MA, USA) and RNA was separated to aqueous phase using chloroform. The aqueous layer containing RNA was then mixed with isopropanol and loaded to Aurum Total RNA Mini Kit columns (Bio-Rad, Mississauga, ON, Canada). Total RNA was then extracted according to the manufacturer’s instructions and concentration was assessed on the nanodrop attachment for the Varioskan LUX Multimode Microplate reader (Thermo Scientific). Reverse transcription of RNA into cDNA was performed by M-MLV reverse transcriptase and oligo(dT) primers (Qiagen, Toronto, ON, Canada). cDNA was then amplified in a CFX384 Touch Real-Time PCR Detection Systems (Bio-Rad) with a SYBR Green master mix and specific primers, listed below in Table 1. Standard curves were created by plotting log DNA concentrations against Ct values from a 1:5 serial dilutions with CFX Manager Software. Gene expression was normalized to a *<u>Rplp0</u>* control ^30^, and relative differences were determined using the ΔΔCt method ^31^. Values are presented as fold increases relative to WT group.

**Table 1.0.**
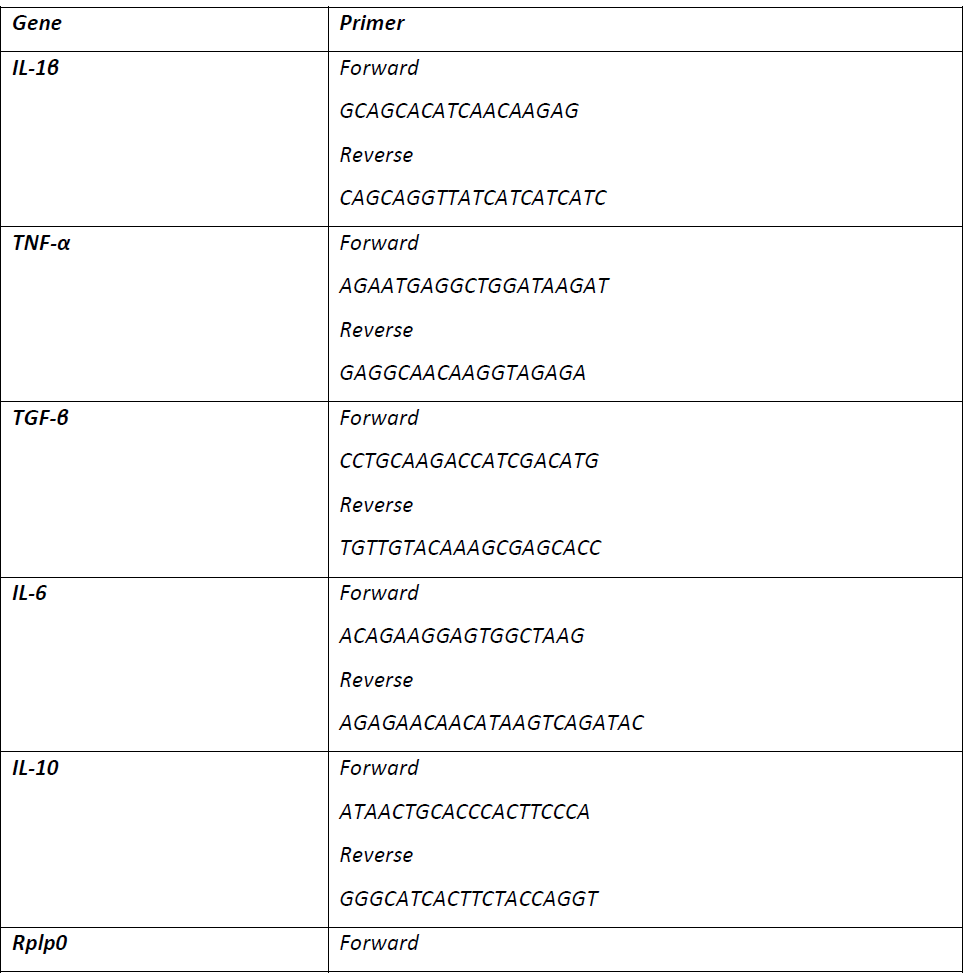
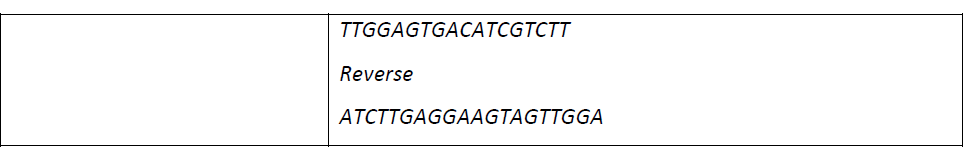
Primers for Cytokine (listed 5’ to 3’)

### Cytokine profiling

Tibialis anterior and diaphragm samples were homogenized in lysis buffer containing (in mM) (20 Tris/HCl, 150 NaCl, 1 EDTA, 1 EGTA, 1% Triton X-100, 2.5 Na_4_O_7_P_2_, 1 Na_3_VO_4_, pH 7.0) supplemented with protease (Roche, Mississauga, ON, Canada) and phosphatase inhibitors (Sigma Aldrich; St. Louis, MI, USA). Samples were diluted two-fold in the assay buffer before loading to the plates with beads in a blinded manner. According to the manufacturer’s instructions, levels of TGF-β, TNF-α, IL-1β, IL-6 and IL-10 in the homogenates were measured using LEGENDplexTM Mouse Custom Panel (BioLegend, San Diego, CA, USA). Assay was performed in the Attune NxT flow cytometer (Thermo Fisher). The FCS files generated on the flow cytometer were analyzed using the LEGENDplex™ cloud-based analysis software.

### Functional assays and serum creatine kinase

Voluntary wheel running, cage hang time and forelimb grip strength were assessed two days before sacrifice and as described previously ^27, 32^. Serum derived from cardiac puncture at the time of sacrifice were assessed spectrofluorometrically for creatine kinase activity (U/L) as described in ^27^.

### In-vitro diaphragm force

Assessment of *in-vitro* diaphragm force was partially adapted from ^33, 34^. The entire diaphragm was excised, placed in ice cold Ringer solution and a strip not exceeding 3-4mm in width was cut as described previously ^33^. Briefly, a silk suture was attached to the central tendons and the ribs and secured in a bath with oxygenated Ringers solution containing (in mM) 121 NaCl, 5 KCl, 1.8 CaCl_2_, 0.5 MgCl_2_, 0.4 NaHPO_4_, 24 NaHCO_3_, 5.5 glucose and 0.1 EDTA; pH 7.3 (95% O_2_/5% CO_2_) maintained at 25°C with ribs secured to the force transducer with an s-hook and central tendon to the lever arm. The diaphragm was positioned between two platinum electrodes driven by a biphasic stimulator (Model 305C; Aurora Scientific, Aurora, ON, Canada), acclimatized for 30 minutes and optimal resting length (L_0_) determined. A force-frequency curve was performed (1, 10, 20, 40, 60, 80, 100, 120, 140, 200 Hz) with 1 minute of rest between contractions with a final 5-minute rest period provided before a fatiguing protocol was performed (70Hz for 350ms every 2 seconds for 5 minutes). After the force-frequency protocol was assessed, a 5 minute rest period was provided before a fatiguing protocol was performed (70Hz for 350ms every 2 seconds for 5 minutes). Recovery from fatigue was assessed at 5-, 10- and 15-minutes post fatigue at the frequency that elicited maximum force in the force-frequency protocol. Force production was analyzed using Dynamic Muscle Control Data Acquisition software (Aurora Scientific) and normalized to cross sectional area (CSA) of the muscle strip (m/l*d) where ‘m’ is the mass of the strip devoid of the central tendon and ribs, ‘l’ is the length (from the point of insert on the ribs to the myotendinous junction of the central tendon) and ‘d’ is the mammalian skeletal muscle density (1.06g/mm^3^) ^35^.

### In-situ quadriceps force

Quadriceps force protocol was adapted from previous work ^36^. Mice were anaesthetized with isoflurane vaporized in medical air (21% oxygen) at a 2L/min flow rate and placed on a platform. Body temperature was maintained using an overhead heat lamp. Once fully sedated, hair was removed from the knee and a small vertical incision was made above the knee to expose the patellar tendon. The tendon was then secured with a suture (4/0; Fine scientific instruments, North Vancouver, Canada), with a loop at the end. The tendon was then severed, and the loop was then attached to an Aurora Scientific 305C muscle lever arm with an s-hook (Aurora Scientific). A 27 G needle was pierced from the lateral side of the knee through the femoral epicondyles immobilizing the knee joint. The knee was then secured with a vertical knee clamp and quadriceps contraction was facilitated through percutaneous stimulation of the femoral nerve. Optimal resting length (L_0_) was determined using single twitches (pulse width=0.2ms) at varying muscle lengths at the maximal stimulation current. Force was assessed across increasing frequencies (1, 40, 60, 80, 100, 120 Hz) using a biphasic stimulator (Model 710B, Aurora Scientific), with one minute rest between each frequency. After a five minute rest period, a fatigue protocol (70Hz stimulation once every 1.5s for 120 contractions) was initiated. Recovery from fatigue was assessed 5, 10 and 15 minutes after the last contraction of the fatigue protocol at the frequency that elicited max force. Data were analyzed using Dynamic Muscle Control Data Acquisition software (Aurora Scientific) and normalized to quadriceps weight in mg.

### Preparation of permeabilized muscle fibre bundles

This technique was adapted from previous methods ^27, 37^. Following completion of *in situ* quadriceps force in one limb, the quadriceps from the non-stimulated limb and diaphragm were excised carefully from mice while under heavy sedation with isoflurane and placed immediately into ice-cold BIOPS buffer. Tibialis anterior was not used due to limitations in tissue availability. The quadriceps from the non-stimulated limb and the diaphragm were collected and quickly placed in a buffer containing (in mM) 50 MES Hydrate, 7.23 K_2_EGTA, 2.77 CaK_2_EGTA, 20 imidazole, 0.5 dithiothreitol, 20 taurine, 5.77 ATP, 15 PCr, and 6.56 MgCl_2_·6 H_2_O (pH 7.1). Both muscles were trimmed of connective tissue and fat and separated along the longitudinal axis into small bundles weighing approximately 1.0-2.5mg wet weight for respiration and mH_2_O_2_ and 0.5-1.5mg wet weight for calcium retention capacity. Bundles were permeabilized with 40µg/µL saponin (Sigma Aldrich) in BIOPS on a platform rotor for 30 minutes at 4°C. The permeabilized bundles were then transferred to wash buffers to remove saponin as follows: MiRO5 for mitochondrial respiration, Buffer Z for mitochondrial H_2_O_2_ emission (mH_2_O_2_), and Buffer Y for calcium retention capacity. The composition of each buffer has been described previously ^27^. Fibres prepared for pyruvate-supported mitochondrial H_2_O_2_ emission (mH_2_O_2_) were permeabilized in the presence of 35µM 2,4-dinitrochlorobenzene (CDNB) to deplete glutathione and enhance the detection of mH_2_O_2_ ^38^. All bioenergetic assays were performed within 4 hours of washing to maintain fibre viability.

### Mitochondrial respiration

Mitochondrial respiration was assessed in permeabilized muscle fibres using high resolution oxygen consumption as an index of mitochondrial respiration. Assessments were performed in 2mL of MiRO5 supplemented with (creatine-dependent) or without (creatine independent) 20mM creatine to saturate mitochondrial creatine kinase activity ^37, 39^. O_2_ consumption was measured using the Oroboros Oxygraph-2K (Oroboros Instruments, Corp., Innsbruck, Austria) while stirring at 37°C in the presence of 5µM blebbistatin to prevent muscle fibre contraction by rigor in response to ADP ^37^. Each chamber was oxygenated with 100% pure O_2_ to an initial concentration of ∼350 μM and experiments were completed before chamber [O_2_] reached 150 μM ^37, 40–43^. Prior to permeabilization, fibre bundles were gently and quickly blotted dry, weighed in ∼1.5 mL of tared cold BIOPS (ATP-containing relaxing media) to ensure fibres remained relaxed. Respiration was normalized to bundle wet weight. Complex I-supported respiration was stimulated using 5mM pyruvate and 2mM malate (NADH; State II respiration) followed by a titration of ADP concentrations (State III) from physiological ranges (25, 100µM) to supraphysiological (500 µM) and saturating to stimulate maximal coupled respiration (5000 µM). An addition of 10µM cytochrome *c* was added to test mitochondrial outer membrane integrity. Experiments with very low ADP-stimulated respiration and high cytochrome c responses were removed. Lastly, 10mM succinate (FADH_2_) was added to support complex-II respiration.

### Mitochondrial H_2_O_2_ emission (mH_2_O_2_)

mH_2_O_2_ emission was assessed in permeabilized muscle fibres as described previously ^27^. Briefly, mH_2_O_2_ was determined spectrofluorometrically (QuantaMaster 40, HORIBA Scientific, Irvine, CA, USA) utilizing Buffer Z containing 10µM Amplex Ultra Red (Life Technologies; Carlsbad, CA, USA), 1U/mL horseradish peroxidase, 1mM EGTA, 40U/mL Cu/Zn SOD1, 5µM blebbistatin and 20mM Cr. Complex I-supported mH_2_O_2_ was initiated through the addition of 10mM pyruvate and 2mM malate (NADH; forward electron flow) and separately with 10mM succinate (FADH_2_; reverse electron flow from Complex II to I). The ability of ADP to suppress mH_2_O_2_ was assessed with a titration of physiological concentrations (25, 100) and saturating for oxidant generation (500µM). All protocols were repeated without creatine in the assay buffer to permit comparisons with the creatine condition, thereby allowing assessments of mitochondrial creatine sensitivity. Bundles were lyophilized in a freeze-dryer (Labconco, Kansas City, MO, USA) for >4 h and weighed on a microbalance (Sartorius Cubis Microbalance, Gottingen, Germany). The rate of mH_2_O_2_ emission was calculated using a standard curve under the same assay conditions and then normalized to fibre bundle dry weight.

### Mitochondrial calcium retention capacity

Mitochondrial calcium retention capacity was assessed spectrofluorometrically as previously described but with modifications ^27, 44^. This assay was measured spectrofluorometrically (QuantaMaster 80, HORIBA Scientific) in a cuvette with 300µL assay buffer containing 1 μM Calcium Green-5N (Invitrogen), 2 μM thapsigargin, 5 μM blebbistatin, and 40 μM EGTA while maintained at 37°C with continuous stirring. 5mM glutamate, 2mM malate, 5mM ADP and 20mM creatine were added to the assay buffer and minimum fluorescence was recorded. 4nmol pulses of CaCl_2_ were added until the mitochondrial permeability transition pore (mPTP) opening was observed as an increase in fluorescence corresponding to net mitochondrial calcium release, at which point saturating pulses of CaCl_2_ were used to establish maximum fluorescence. Changes in free Ca^2+^ during mitochondrial Ca^2+^ uptake were then calculated using the known K_d_ for Calcium Green-5N and equations established for calculating free ion concentration ^45^. Calcium retention was then normalized to fibre bundle dry weight.

### Histochemical & immunofluorescent staining

Excised muscles were mounted for histology and immunofluorescence in Tissue Plus Optimal Cutting Temperature compound (OCT; Fisher Healthcare), frozen in isopentane chilled in liquid nitrogen and stored at −80°C. Samples collected were cut into 8µm thick serial cross sections with a cryostat (Thermo Fisher Scientific; Kalamazoo, MI, United States;) maintained at −20°C on Fisherbrand Superfrost Plus slides (Thermo Fisher Scientific). <u>All images were assigned arbitrary numbers and examined without</u> <u>knowledge of the treatment or group (i.e., blinded analysis).</u> Hematoxylin and eosin (H&E) and Masson’s trichrome staining protocols are described elsewhere ^27^. H&E sections images were taken using EVOS M7000 imager (Thermo Fisher Scientific) using 20X magnification and analyzed using ImageJ (http://imagej.nih.gov/ij/). for areas of damage (fibres with fragmented sarcoplasm and immune cell infiltration) and expressed as a percentage of the entire muscle section. To determine fibrotic area, skeletal muscle sections were stained with Masson’s Trichrome (fibrotic regions stain blue for collagen). Sections were imaged Nikon 90i-eclipse microscope (Nikon Inc., Melville, NY, USA) and analyzed with the NIS Elements AR software (v4.6, Nikon). Immunofluorescent analysis of myosin heavy chain expression was previously described ^46^. Images were taken with EVOS M7000 equipped with standard red, green, blue filters cubes. Fibres that were not positively stained were considered IIX fibres. These images were then analyzed with ImageJ for minimal Feret’s diameter as an index of muscle atrophy ^47^.

### Western blotting

Frozen sections of quadriceps and diaphragm from each muscle (approximately 10-30mg in size) were homogenized using a plastic microcentrifuge tube with tapered Teflon pestle in ice-cold lysis buffer containing (in mM) (20 Tris/HCl, 150 NaCl, 1 EDTA, 1 EGTA, 1% Triton X-100, 2.5 Na_4_O_7_P_2_, 1 Na_3_VO_4_, pH 7.0) supplemented with protease (Sigma Aldrich) and phosphatase inhibitors (Roche). A 12% gel was used to separate electron transport chain proteins and all other proteins were separated on a 10% gel. Following, all gels were transferred onto 0.2µm low fluorescence PVDF membrane (Bio-Rad, Mississauga, Canada) and blocked with Li-COR Intercept Blocking Buffer (LI-COR, Lincoln NE, USA) for 1 hour. Membranes were then incubated with specific primary antibodies (listed below) overnight at 4°C. A commercially available monoclonal rodent OXPHOS Cocktail (ab110413; Abcam, Cambridge, UK, 1:250 dilution) including V-ATP5A (55 kDa), III-UQCRC2 (48 kDa), IV-MTCO1 (40 kDa), II-SDHB (30 kDa), and I-NDUFB8 (20 kDa) were used to detect electron transport chain proteins. Commercially available polyclonal antibodies were used to detect p-AMPKα (Thr172; 62kDa) (2535, Cell Signaling Technologies (CST), Danvers, MA, USA; 1:1000), AMPKα (62 kDa; 2532, CST, 1:500), p-p-38MAPK (Thr180/Tyr182;43 kDa;) (9211, CST, 1:1000) and p38MAPK (40 kDa; 9212, CST, 1:500). After overnight primary antibody incubation, membranes were washed three times (5 minutes each time) in TBS-T and incubated at room temperature with appropriate fluorescent secondary antibody (LI-COR). Prior to detection, membranes were washed in TBS-T three times for 5 minutes and then imaged using infrared imaging (LI-COR CLx; LI-COR) and quantified by densitometry (ImageJ, http://imagej.nih.gov/ij/). All images were normalized total protein from the same membrane stained using Amido Black total protein-stained membrane (A8181, Sigma).

### Statistical analyses

Results are expressed as means ± SD with the level of significance established as *P* < 0.05 for all statistics. Prior to statistical analyses, outliers were omitted in accordance with ROUT testing (Q=0.5%) and then tested for normality using a D’Agostino–Pearson omnibus normality test (GraphPad Prism Software, La Jolla, CA, USA). State III respiration and mH_2_O_2_ were analyzed using a two-way ANOVA followed by a two-stage step-up method of Benjamini, Krieger and Yekutieli for controlling False Discovery rate (FDR) for multiple-group comparisons. For all other data, statistical differences were analyzed using a one-way ANOVA between all groups followed by a two-stage step-up method of Benjamini, Krieger and Yekutieli post hoc analyses where appropriate. All reported p-values are FDR-adjusted p values (traditionally termed ‘q’).

## Results

### ALY688-SR treatment attenuates fibrosis in the diaphragm of D2.mdx mice

We first examined the effect of ALY688-SR treatment on diaphragm fibrosis defined as the absolute increase in collagen relative to levels observed in wildtype (WT). Using Masson’s trichrome staining (Figure 1B), a 5.5-fold increase in collagen content was observed in diaphragm from vehicle treated D2.*mdx* mice indicating fibrosis is present at a very early stage of disease development (VEH vs WT, Figure 1B). Remarkably, both doses of ALY688-SR attenuated this fibrosis by 66.7-67.6%. Collagen content increased from 0.5% to 1.1% in tibialis anterior in D2.*mdx*-VEH mice compared to WT (Figure 1C) which indicates the diaphragm is more severely affected at this young age. Consequently, there was no detectable effect of either dose of the drug. Collectively, these findings indicate that early treatment with ALY688 decreases fibrosis in muscles severely affected by dystrophin mutation.

**Figure 1:**
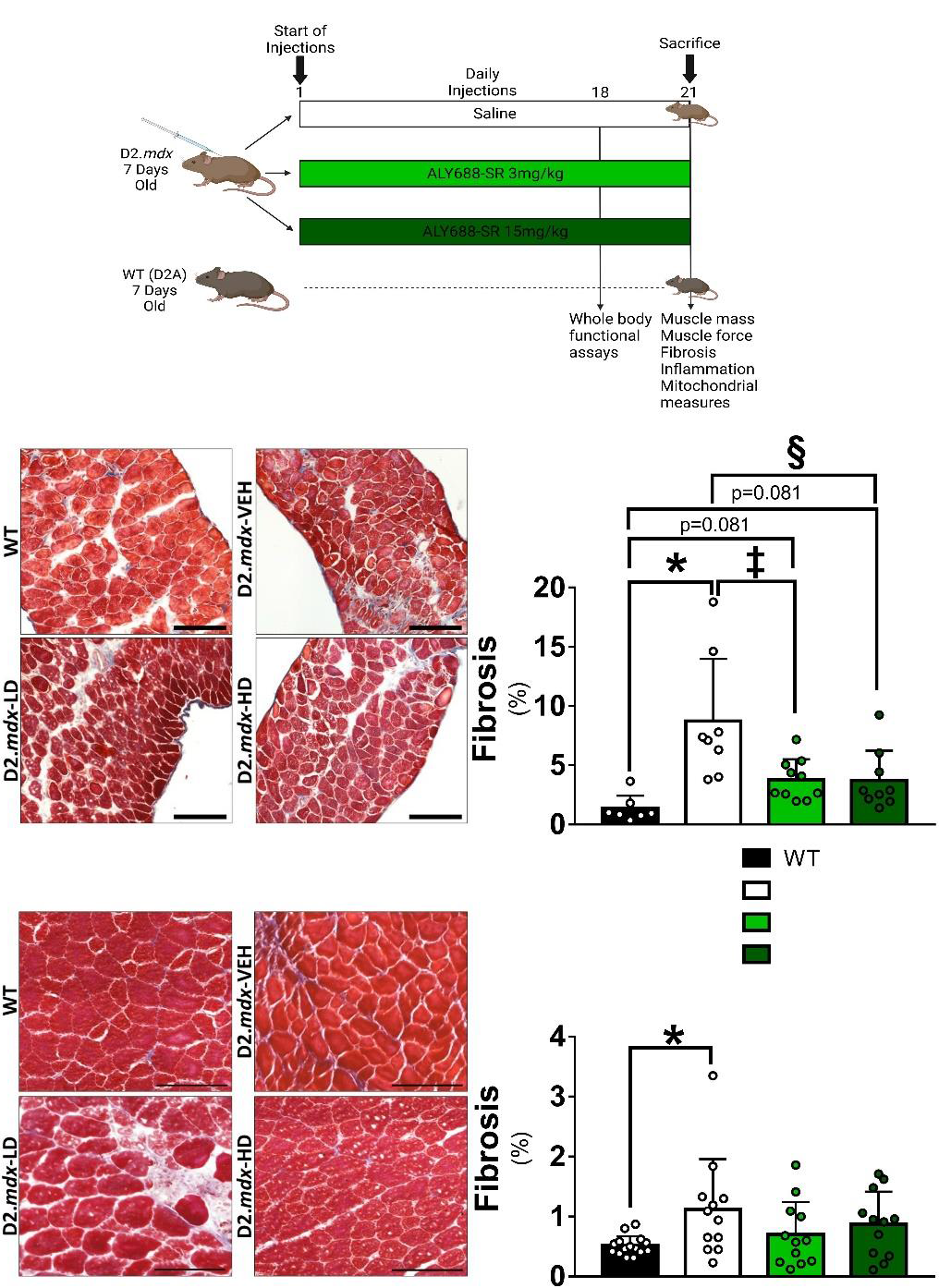
Schematic of study design and quantification of interfibrillar collagen accumulation in D2.*mdx* mice treated with ALY688-SR. A visual representation of study design is provided in (A). 7 day old D2.*mdx* mice received subcutaneous injections of ALY688-SR at a low dose (LD; 3mg/kg body weight) or high dose (HD; 15mg/kg body weight) or a saline control for 21 days. Healthy D2A mice (wild type (WT)) were used as age matched controls. On day 18 of treatment, all four groups underwent whole body functional assays to assess voluntary muscle function. After 21 days of injections, all groups were sacrificed, and tissue collected for the outlined measures. Fibrosis was evaluated in diaphragm (B) and tibialis anterior (C) using Masson’s trichrome staining. Results represent mean ±SD; n=7-15. All p values are FDR-adjusted by Benjamini, Krieger and Yekutieli post-hoc analyses. *p<0.05 WT vs D2.*mdx*-VEH; ‡p<0.05 D2.*mdx*-VEH vs D2.*mdx*-LD; §p<0.05 D2.*mdx*-VEH vs D2.*mdx*-HD. Representative images of diaphragm (B, magnification, ×20) and tibialis anterior (C, magnification, ×20). Scale bar =100µm.

### ALY688-SR attenuates atrophy of D2.mdx diaphragm in a fibre type-specific manner

We also examined minimal feret diameter in each fibre type as a measure of muscle atrophy. The high dose of ALY688-SR completely prevented atrophy in myosin heavy chain (MHC) I fibres. Additionally, in IIA fibres, minimal feret diameter was reduced from 24µm in WT to 20µm in VEH compared to 22µm in HD (Figure 2A). In IIB fibres, minimal feret diameter was reduced from 26µm in WT to 23µm in VEH compared to 24µm in HD. A similar prevention of atrophy in IIB fibres was seen in LD.

**Figure 2:**
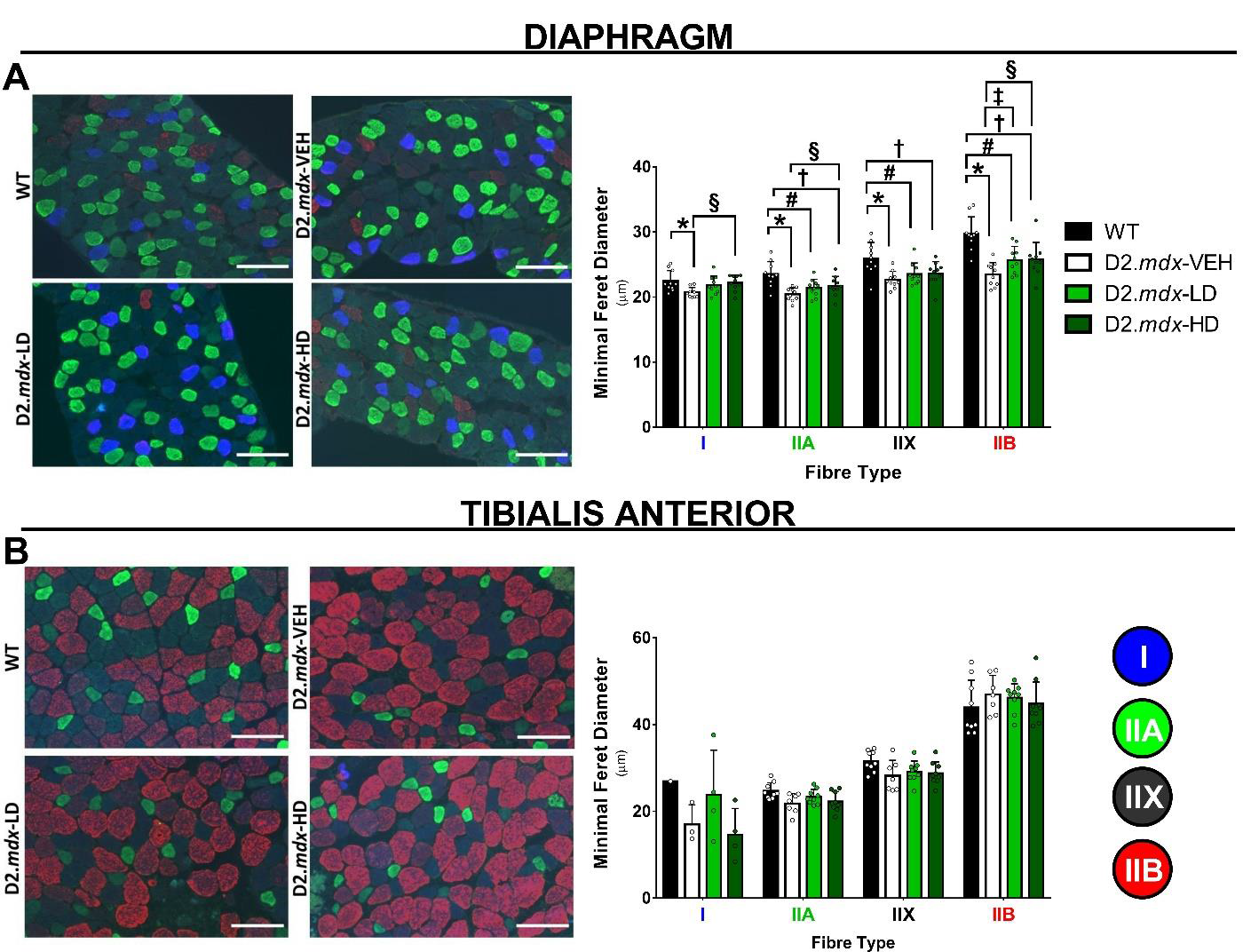
Assessments of fibre-type specific minimal feret diameter in D2.*mdx* mice treated with ALY688-SR. Minimal feret diameter was assessed in diaphragm (A) and tibialis anterior (B) using immunofluorescent detection of myosin heavy chain isoforms. Results represent mean ±SD; n=7-10. All p values are FDR-adjusted by Benjamini, Krieger and Yekutieli post-hoc analyses. *p<0.05 WT vs D2.*mdx*-VEH; #p<0.05 WT vs D2.*mdx-* LD; †p<0.05 WT vs D2.*mdx-* HD; ‡p<0.05 D2.*mdx*-VEH vs D2.*mdx*-LD; §p<0.05 D2.*mdx*-VEH vs D2.*mdx*-HD. Representative images of diaphragm (A, magnification, ×20) and tibialis anterior (C, magnification, ×20). Scale bar =100µm.

There were no differences in minimal feret diameter in any fibre type of tibialis anterior between the groups (Figure 2B). This finding suggests that the modest fibrosis in tibialis anterior seen at a young age (Figure 1C) precedes atrophy given previous studies have shown reductions in fibre size in this muscle at later stages of disease in *mdx* mice ^48^.

### ALY688-SR treatment alters markers of inflammation

We next examined the effect of ALY688-SR on markers of muscle damage and inflammation in the diaphragm and tibialis anterior. H&E staining was used to assess the degree of damaged areas consisting of areas of necrosis that often correspond to fibrosis (Figure 3A and Supplemental Figure 1A). All D2.*mdx* groups demonstrated an increase in damaged areas (3.7-to-3.9-fold change vs WT; Figure 3A) suggesting the attenuated fibrosis (Figure 1B) with the drug occurred independently of generalized fibre damage. Next, we chose to assess inflammation only with high dose ALY688-SR given that both doses were equally as effective in attenuating fibrosis but the high dose ALY688-SR was more consistent in preventing atrophy. IL-6 mRNA content was increased in diaphragm from D2.*mdx*-VEH mice which was attenuated by the high dose (51.8% increase vs WT; Figure 3B). In contrast, ALY688-SR increased IL-6 protein content in the diaphragm (69.1% increase vs VEH; Figure 3C). This result in muscle tissue lysate does not rule out the possibility that expression of IL-6 in non-muscle cell types occurred (see Discussion). Increases in IL-1β mRNA in D2.*mdx*-VEH mice (2.7-to-3.0-fold increase vs WT) were not affected by the drug (Figure 3B) while protein content was similar between all groups (Figure 3C). TNF-α and IL-10 mRNA and protein contents were not different between groups (Figure 3). TGF-β protein content was increased in response to the drug (20.3% increase vs VEH; Figure 3C) although mRNA contents were unchanged in all groups (Figure 3B).

**Figure 3:**
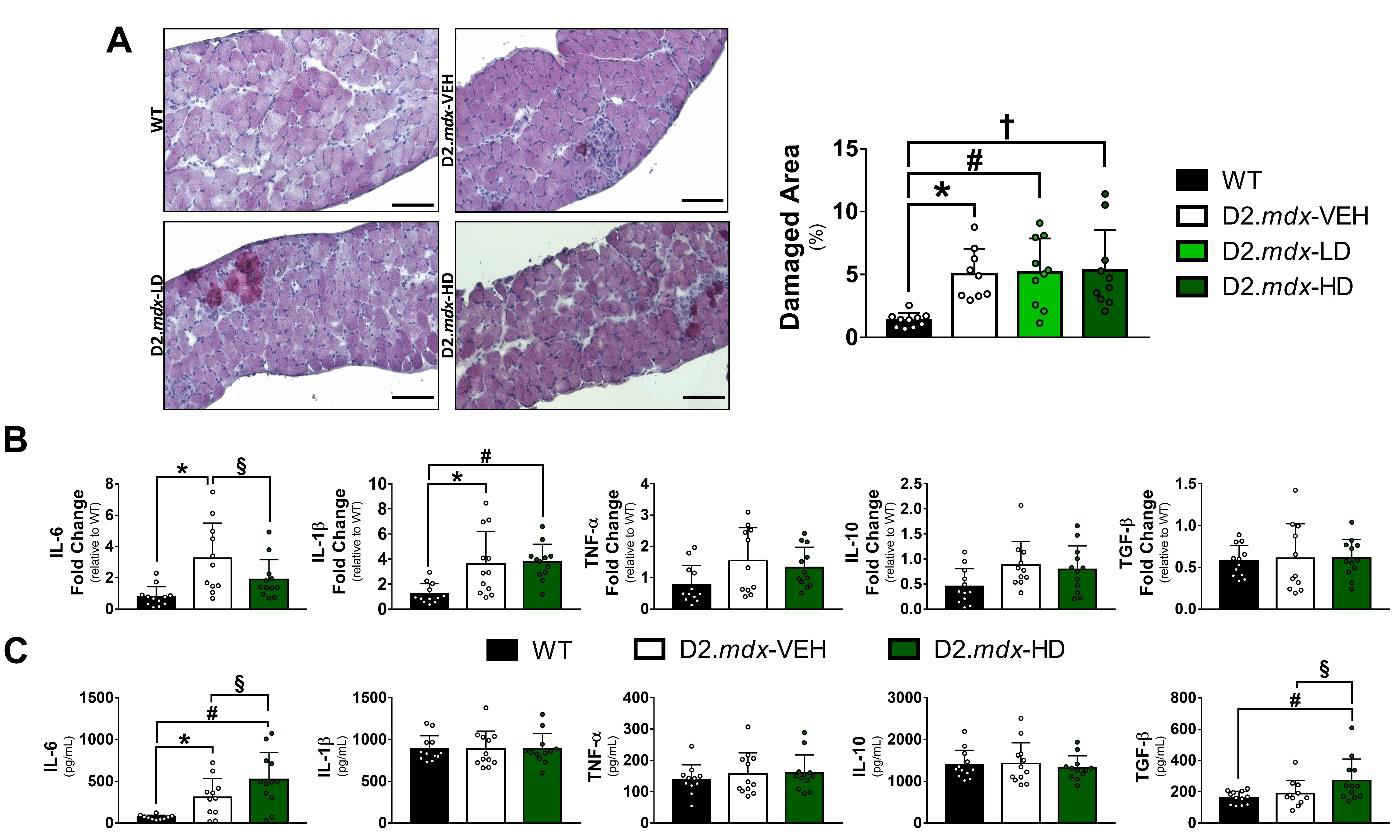
Effects of ALY688-SR on inflammation and muscle damage in D2.*mdx* diaphragm. As both doses of ALY688 were effective in reducing fibrosis in diaphragm, only high dose groups were examined in inflammation. (A) Hematoxylin & eosin staining were used to assess areas of muscle damage which includes areas of necrosis and fibrosis and expressed as a percentage of total area. Scale bar =100µm. (B) qPCR were used to assess mRNA fold changes of IL-6, IL-1β, TNF-α, IL-10 and TGF-β across WT and D2.*mdx* mice treated with vehicle or high dose ALY688-SR treatment. qpCR results were normalized to *Rplp0*, and the subsequent ratios were presented as relative expression compared with WT values. (C) Protein levels of the same cytokines were assessed by BioLegend Multiplex using flow cytometry. Results represent mean ±SD; n=7-12. All p values are FDR-adjusted by Benjamini, Krieger and Yekutieli post-hoc analyses. *p<0.05 WT vs D2.*mdx*-VEH; #p<0.05 WT vs D2.*mdx-* LD; †p<0.05 WT vs D2.*mdx-* HD; §p<0.05 D2.*mdx*-VEH vs D2.*mdx*-HD.

In the tibialis anterior, robust increases in damaged areas were noted in all three D2.*mdx* groups (17.2-to-22.6-fold increase vs WT; Supplemental Figure 1A) in contrast to the small degree of fibrosis seen in Figure 1. D2.*mdx*-VEH mice demonstrated greater mRNA contents of TNF-α, IL-1β, IL-6, IL-10 and TGF-β which were paralleled by greater protein contents for TGF-β (9.7-fold increase vs VEH; Supplemental Figure 2B, C) compared to wildtype. The high dose of ALY688-SR attenuated TNF-α mRNA compared to VEH (41.4%; Supplemental Figure 1B). TNF-α protein content was not detectable in wildtype but increased to 40.6 pg/ml in VEH. HD prevented 83.9% of this increase (Supplemental Figure 1C).

### ALY688-SR lowers mitochondrial H_2_O_2_ emission by enhancing ADP responsiveness

Mitochondria produce H_2_O_2_ during oxidative phosphorylation of ADP to ATP. As cellular consumption of ATP increases, the cycling of ADP to the matrix is accelerated. Increasing matrix ADP attenuates membrane potential which in turn lowers mH_2_O_2_ emission given the two latter processes are inversely proportional ^49^. By considering these principles, we designed *in vitro* assay conditions to capture the effect of increasing metabolic demand on oxidant generation by titrating ADP following induction of mH_2_O_2_ emission from mitochondria with NADH-generating substrates (pyruvate/malate) to support Complex I (Figure 4A). This approach revealed that mH_2_O_2_ emission was higher at each ADP concentration in D2.*mdx* diaphragm and quadriceps (2.0-to-9.1-fold increase vs WT; Figure 4B, C). This elevation was due specifically to a reduced ability of ADP to attenuate mH_2_O_2_ relative to wildtype muscle which is similar to our previous findings ^27, 28, 50, 51^. The results suggest that dystrophin mutation leads to a greater degree of mH_2_O_2_ emission across a range of oxidative phosphorylation kinetics. The high dose of ALY688-SR completely preserved mitochondrial responsiveness to ADP and restored mH_2_O_2_ emission to normal kinetics in diaphragm (8.7 to 70.7%; Figure 4B). In quadriceps, both doses of the drug partially attenuated mH_2_O_2_ emission (45.9 to 62.3%; Figure 4C). mH_2_O_2_ emission was not assessed in tibialis anterior.

**Figure 4:**
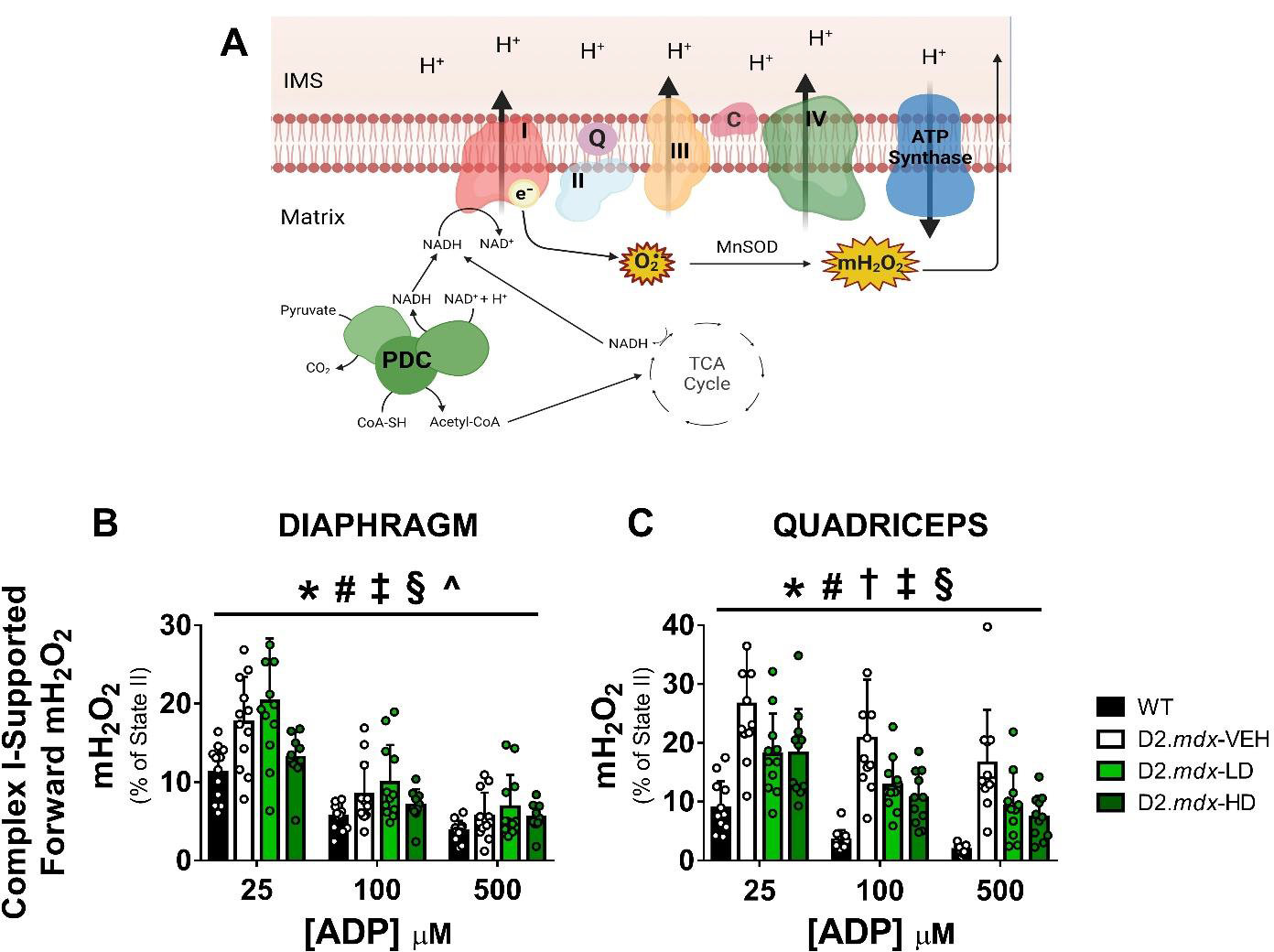
ALY688-SR enhances ADP-suppression of complex-I supported mitochondrial H_2_O_2_ emission. (A) Schematic representation of complex-I supported mitochondrial H_2_O_2_ emission (mH_2_O_2_) due to forward electron flow from NADH generated by pyruvate. In states of high membrane potential (low ADP), electrons at complex I may ‘slip’ prematurely at, for example, Complex I to produce superoxide radicals (O_2_^•-^) which are then converted to H_2_O_2_ by manganese superoxide dismutase (MnSOD). 10mM pyruvate and 2mM malate (to assist with continual flux through pyruvate dehydrogenase complex (PDC)), were used to stimulate electron flow through complex I in permeabilized fibre bundles from diaphragm (B) and quadriceps (C) in the absence of ADP (state II; high membrane potential). Subsequent titrations of physiological levels of ADP (‘resting muscle’, 25uM; high periods of demand similar to intense exercise; 100uM) and supramaximal (500uM) attenuates mH_2_O_2_ by promoting forward electron flow and is expressed as a % of State II mH_2_O_2_ emission to reveal the degree of attenuation in response to ADP ^39, 87, 88^. Results represent mean ±SD; n=10-12. All p values are FDR-adjusted by Benjamini, Krieger and Yekutieli post-hoc analyses. *p<0.05 WT vs D2.*mdx*-VEH; #p<0.05 WT vs D2.*mdx-* LD; †p<0.05 WT vs D2.*mdx-* HD; ‡p<0.05 D2.*mdx*-VEH vs D2.*mdx*-LD; §p<0.05 D2.*mdx*-VEH vs D2.*mdx*-HD; ^p<0.05 D2.*mdx*-LD vs D2.*mdx*-HD. TCA= tricarboxylic cycle; PDC= pyruvate dehydrogenase complex. O_2_^•-^= superoxide radicals

In separate *in vitro* experiments, we also used succinate (FADH_2_; Complex II) to stimulate oxidant generation at Complex I as occurs through a unique reverse electron flow from Complex II to Complex I ^52, 53^. Furthermore, we compared these kinetics in both the absence and presence of creatine given creatine accelerates matrix ADP/ATP cycling which can alter the influence of ADP in attenuating mH_2_O_2_, at least in response to succinate ^54^. Using this approach, we found that succinate-induced mH_2_O_2_ emission was also elevated in D2.*mdx* diaphragm but there was no effect of either drug dose (1.1-to-2.3-fold increase vs WT; Supplemental Figure 2A, C). However, increased mH_2_O_2_ emission seen in quadriceps from D2.*mdx* mice were robustly attenuated by both doses (11.3 to 73.6%; Supplemental Figure 2B, D).

Thus, while high doses lowered mitochondrial H_2_O_2_ emission due to forward electron flow from Complex I and supported by glucose-derived substrate (pyruvate) in both muscles, succinate-induced mH_2_O_2_ emission by reverse electron flow to Complex I was attenuated only in the quadriceps. These findings highlight the complexity of mitochondrial redox regulation across muscle types, the overall findings demonstrate that ALY688-SR lowers mH_2_O_2_ by preserving mitochondrial responsiveness to ADP.

We also assessed mitochondrial respiration as an index of oxidative phosphorylation in both quadriceps and diaphragm. D2.*mdx* mice demonstrated reduced ADP-stimulated respiration in multiple substrate conditions with generally no effect of either dose in both muscles (−25.9 to −67.0% vs WT; Figure 5A, B; Supplemental Figure 3A-D; Supplemental Table 2). Assessments of mitochondrial content markers through western blotting of electron transport chain complexes subunits were also completed in both muscles (Figure 5C, D). Overall, specific subunits of complex III and V, were significantly lowered in disease state groups in both the quadriceps and diaphragm (−14.7 to −31.5% vs WT) and total sum of complexes were reduced in diaphragm (−13.8 to −19.3% vs WT). HD treatment significantly reduced total sum in quadriceps only (−29.1% vs WT) while LD treatment significantly lowered complex IV content in the diaphragm (−27.0% vs WT).

**Figure 5:**
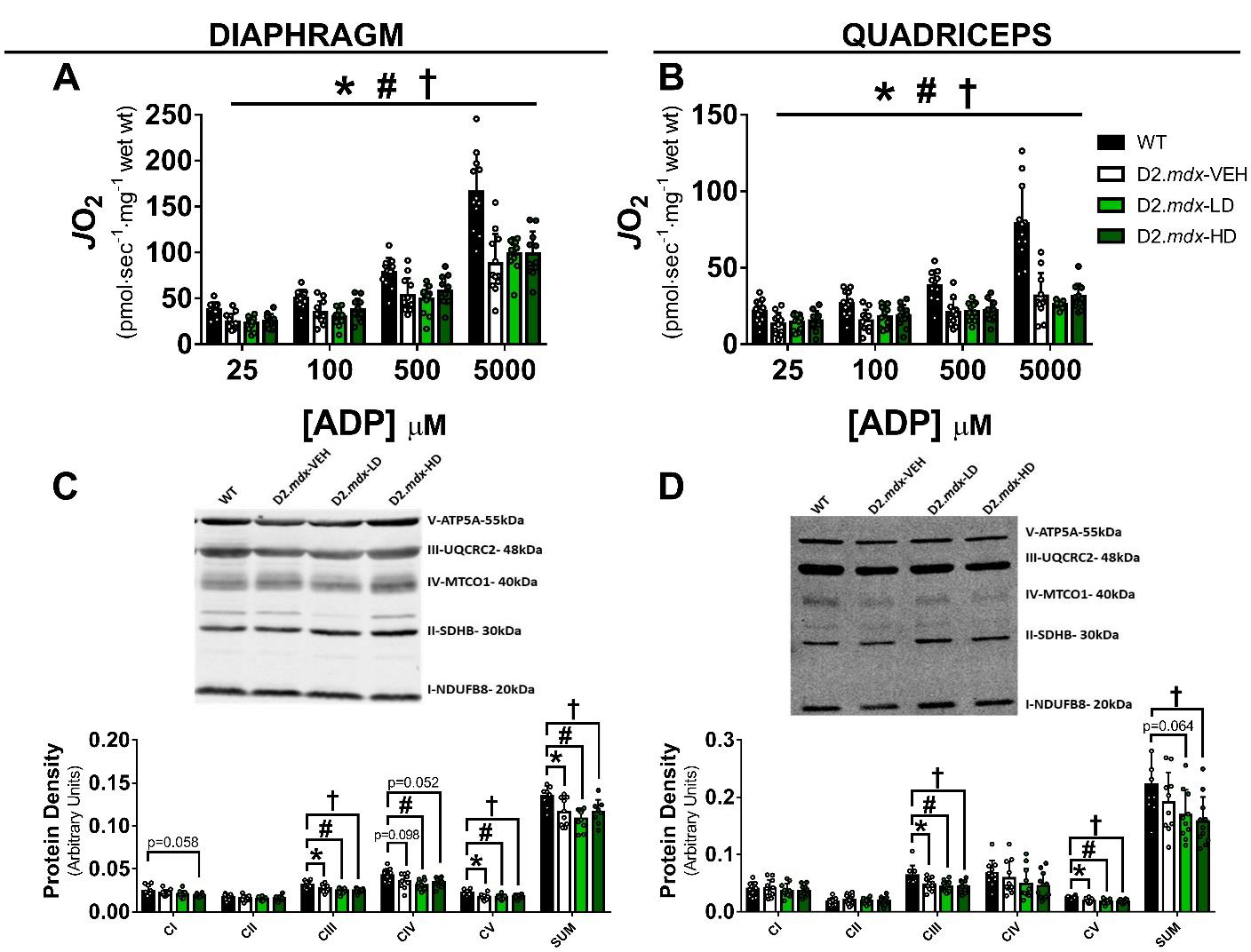
Reductions in pyruvate-supported mitochondrial respiration and protein markers of the electron transport chain in D2.*mdx* mice are not altered by ALY688-SR. Complex-I supported respiration (assessed by oxygen flux, *J*O_2_) stimulated by NADH generation through 5mM pyruvate and 2mM malate in permeabilized fibre across a range of ADP concentrations representing increasing states of metabolic demand: 25μM (modeling resting muscle ^39^); 100μM (modeling high intensity exercise ^88^), 500uM (maximal)^87^ and 5000μM (supramaximal) was assessed in the diaphragm (A) and quadriceps (B). Subunits of complexes I, II, III, IV and V were assessed by western blot in diaphragm (C) and quadriceps (D). Results represent mean ±SD; n=9-12. All p values are FDR-adjusted by Benjamini, Krieger and Yekutieli post-hoc analyses. *p<0.05 WT vs D2.*mdx*-VEH; #p<0.05 WT vs D2.*mdx-* LD; †p<0.05 WT vs D2.*mdx-* HD.

In addition to mH_2_O_2_ emission and respiration, we also assessed the potential for calcium-induced mitochondrial induction of apoptosis given calcium stress occurs in Duchenne muscular dystrophy ^55–57^. However, there was no differences between groups when assessing mitochondrial calcium retention capacity as a marker of permeability transition pore formation (Supplemental Figure 4). Collectively, these mitochondrial assessments point to a more specific relationship between muscle fibrosis, atrophy and mitochondrial redox stress underlying the effects of ALY688-SR on diaphragm in D2.*mdx* mice.

### ALY688-SR remodels muscle force production

We next determined whether muscle force production was different between groups. As expected, D2.*mdx* mice demonstrated lower muscle force production in both diaphragm strips and quadriceps (Figure 6A, B). Both doses of ALY688-SR lowered force relative to both wildtype and D2.*mdx* in diaphragm (−2.3% to –43.0% vs VEH & WT) but had no effect on quadriceps. There was less recovery from fatigue in diaphragm with the high dose and in quadriceps with the low dose (−5.3% to −50.9%; Supplemental Figure 7A, B). While initially unexpected, these assessments did not translate to attenuated physical activity behaviours as noted by similar grip strength, cage hang time and voluntary wheel running between all D2.*mdx* groups (−15.1% to −91.0% vs WT Figure 6C-E) or serum creatine kinase (marker of muscle damage) in the high dose group despite increases seen with the low dose (3.1-to-5.2-fold increase vs WT; Figure 6F).

**Figure 6:**
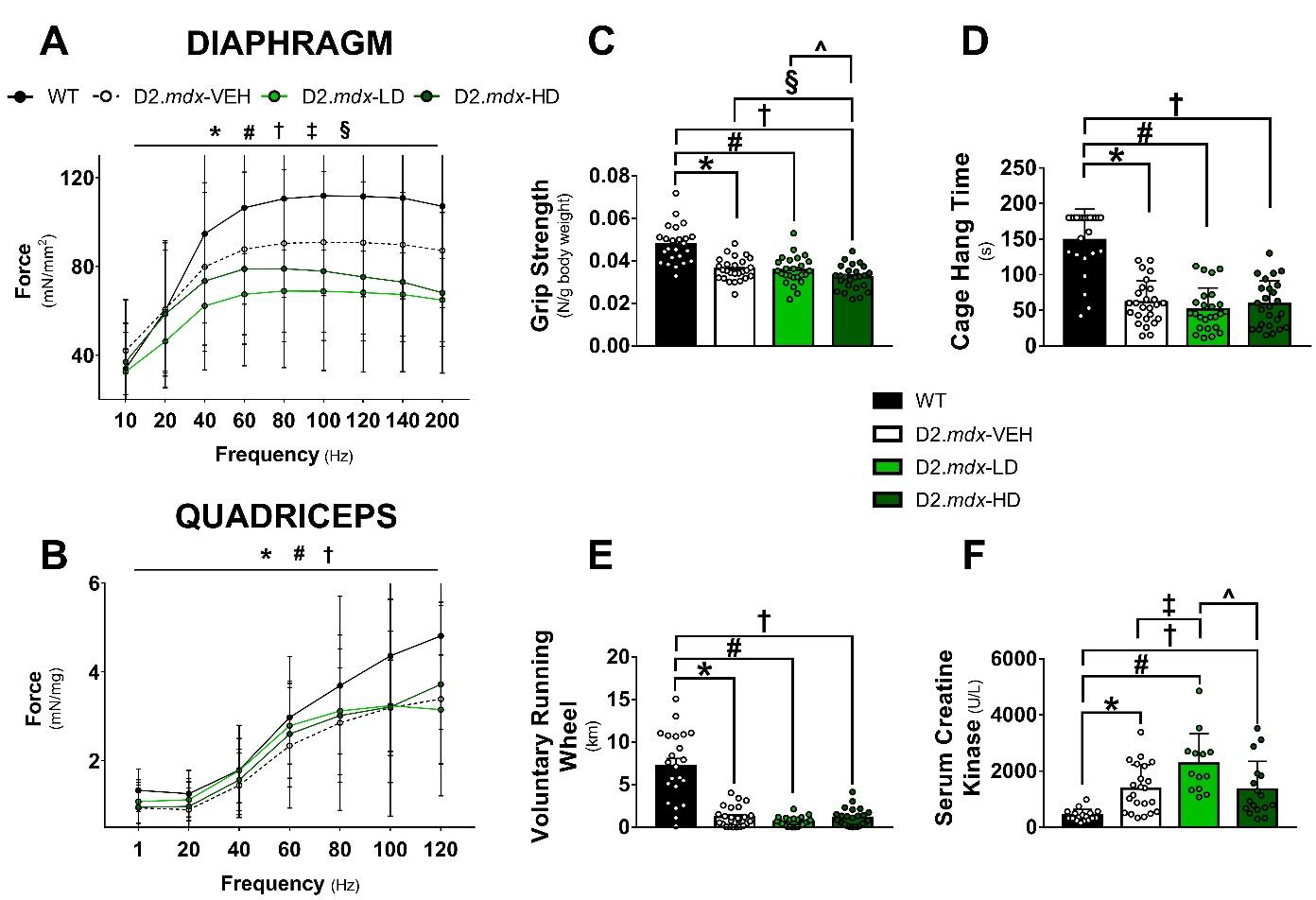
Muscle function assessments of D2.*mdx* treated with ALY688-SR. Muscle force was assessed using force frequency protocols in *in-vitro* diaphragm strips (A) and in-situ quadriceps preparation (B). Serum creatine kinase was assessed as a clinical marker of muscle breakdown (C). Cage hang time (D), grip strength normalized to body weight (E), and distance travelled during 24 hours of voluntary wheel running (F) were assessed as measures of voluntary motor function. Results represent mean ±SD; n=8-13 (A,B); n=13-22 (C); n=21-26 (D-F). All p values are FDR-adjusted by Benjamini, Krieger and Yekutieli post-hoc analyses. *p<0.05 WT vs D2.*mdx*-VEH; #p<0.05 WT vs D2.*mdx-* LD; †p<0.05 WT vs D2.*mdx-* HD; ‡p<0.05 D2.*mdx*-VEH vs D2.*mdx*-LD; §p<0.05 D2.*mdx*-VEH vs D2.*mdx*-HD; ^p<0.05 D2.*mdx*-LD vs D2.*mdx*-HD.

Collectively, these results suggest that the ability of ALY688-SR to attenuate both fibrosis and atrophy in diaphragm are not paralleled by increased force production. However, these force assessments were not compared to broader physiological indicators of respiratory drive and demand that may contribute to adaptive reprogramming of force generating capacity (See Discussion).

### ALY688-SR did not alter AMPK or p38MAPK signaling in D2.mdx mice

ALY688-SR and other adiponectin receptor agonists are thought to attenuate inflammation in part by activating AMPK ^8, 58, 59^. As AMPK is also activated in *mdx* mice ^60^, the potential combined effect of ALY688-SR and disease state in D2.*mdx* mice becomes difficult to predict. Similar to inflammatory measures, only the high dose group was assessed. As shown in Figure 7, AMPK phosphorylation was elevated in D2.*mdx* in both diaphragm and quadriceps (33.2% to 69.1% vs WT). The high dose of ALY688-SR had no effect in either muscle (Figure 7A-C, G-I). As tissues were assessed ∼20-24 hours since the last injection, earlier induction of AMPK may have occurred given prior work showed rapid increases in AMPK phosphorylation in L6 skeletal muscle cells after 30 minutes of treatment ^61^. Phosphorylation of p38 MAPK was also assessed given ALY688-SR has been shown to activate this pathway in previous studies ^62–64^. However, there were no differences in this measure between any of the groups.

**Figure 7:**
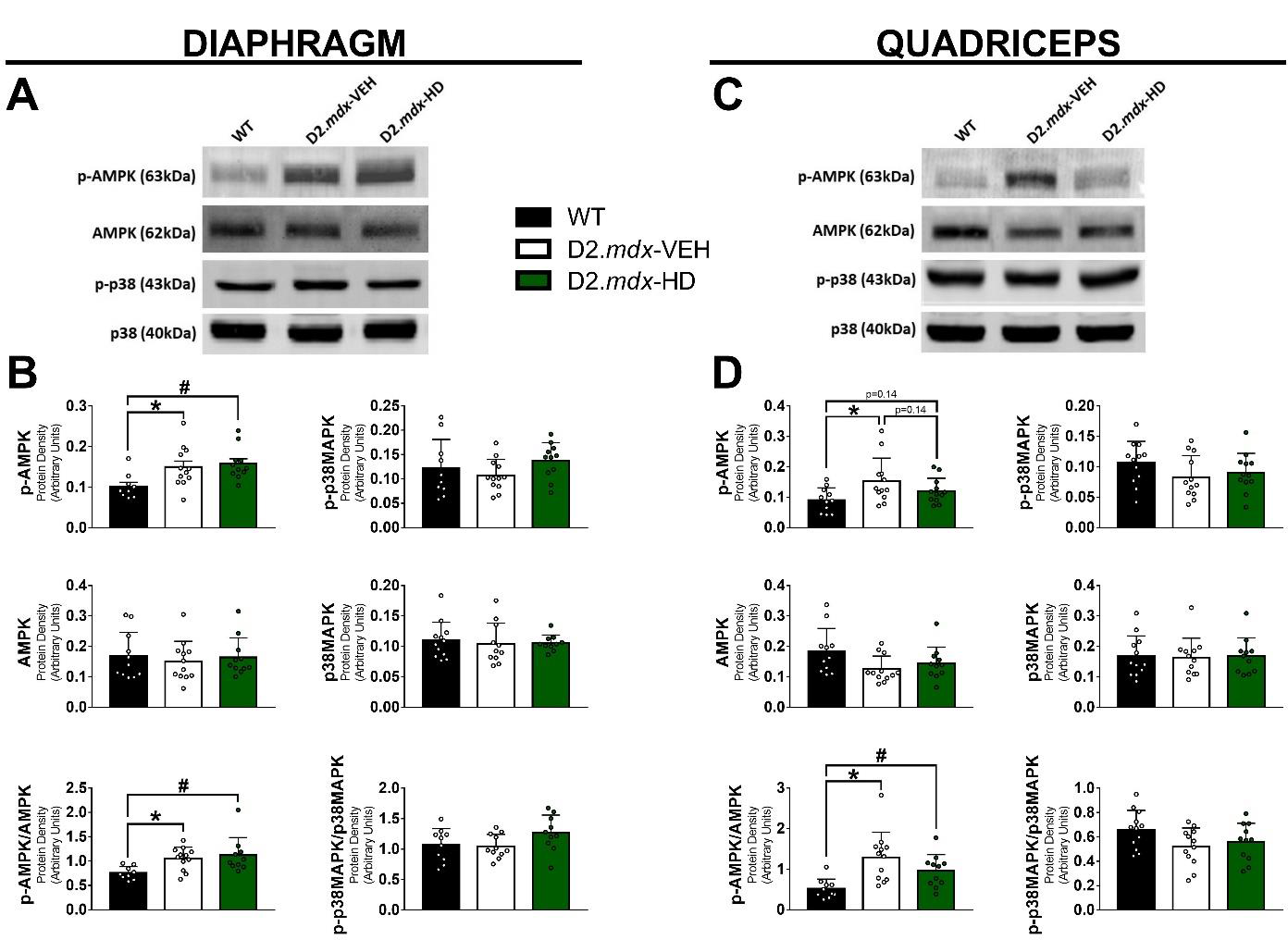
Effects of ALY688-SR treatment on AMPK and p38MAPK activity in D2.*mdx* mice. Western blotting was used to quantify the active phosphorylated form of AMPK, total AMPK, phosphorylated p38MAPK, p38MAPK with normalization of phosphorylated forms to total contents as an index of activity in the diaphragm (A,B) and quadriceps (C,D). Low dose treated animals were excluded in this analysis as high dose treatment attenuated more aspects of dystrophic pathology. Results represent mean ±SD; n=9-12. All p values are FDR-adjusted by Benjamini, Krieger and Yekutieli post-hoc analyses. *p<0.05 WT vs D2.*mdx*-VEH; #p<0.05 WT vs D2.*mdx-* LD.

## Discussion

Progressive fibrosis in diaphragm and limb muscles is believed to contribute to muscle dysfunction in people with DMD and dystrophin-deficient *mdx* mice ^65–67^. While chronic glucocorticoid treatment is associated with delayed age-related increases in fibrosis, at least in heart ^68^, there remains an unmet need to therapeutically prevent fibrosis in addition to preserving muscle mass and function. Here, we identify the adiponectin receptor agonist ALY688-SR as an anti-fibrotic that reduced collagen content in the diaphragm by 66.7-67.6% of very young D2.*mdx* mice after 3 weeks of daily treatment. ALY688-SR also partially attenuated diaphragm atrophy in multiple fibre types. These effects were associated with lower diaphragm mitochondrial reactive oxygen species which suggests a relationship exists between mitochondrial stress and myopathy in this model.

### Disease-modifying effects of ALY688-SR are more apparent in the severely fibrotic diaphragm

The partial prevention of fibrosis and atrophy in the diaphragm was not observed in the tibialis anterior. However, this is likely due to the minimal disease effects noted in this muscle at this young age demonstrating a heterogeneous response of muscle at this early stage of disease progression in the D2.*mdx* mouse. Specifically, when comparing D2.*mdx* vehicle-treated mice to wildtype, collagen content was many-fold higher in diaphragm but only two-fold higher in tibialis anterior. The results indicate that ALY688-SR is effective at partially preventing fibrosis and atrophy when a robust disease state is present in muscle (e.g. diaphragm). Future studies could consider longer term treatment effects of ALY688-SR in muscles that develop more robust fibrosis and atrophy.

These effects in the diaphragm suggest that adiponectin receptor agonism can regulate muscle fibre size and fibrosis. As ALY688 is known to attenuate markers of inflammation ^18^, we determined whether such preservations were related to lower inflammation. While ALY688-SR lowered mRNA content of IL-6, increases in IL-6 protein were also observed compared to vehicle-treated D2.*mdx* mice. This finding is difficult to explain given IL-6 has been shown to be directly associated with fibrosis ^69–73^. While speculative, this divergent response in IL-6 mRNA and protein measured in diaphragm lysate may be related to differential effects of the drug on muscle fibre expression of IL-6 distinct from production in other cell types such as fibroblasts and immune cells. While recombinant adiponectin was shown to increase IL-6 production in human bronchial epithelial cells ^74^, the role of IL-6 in regulating fibrosis is also somewhat controversial given this cytokine is known to have both pro-inflammatory and anti-inflammatory effects depending on the type of experimental model, disease state or systemic stressor^75^. However, mononuclear cell infiltrate assessed by H&E staining was not affected by the drug suggesting changes in fibroblast or immune cell cytokine production may be due to altered activity independent of the cellular contents in muscle tissue.

TGF-β protein content was also increased by ALY688-SR in the diaphragm while mRNA content was not altered, again suggesting a possible increase in production from other cell types within muscle. TGF-β plays an important role in activating fibrosis ^76^, and was previously shown to be induced by the adiponectin receptor ^77, 78^. Additionally, TGF-β is a pleiotropic cytokine that can activate complex signaling pathways with crosstalk at multiple levels leading to highly context-dependent and multifaceted role in health and disease. For example, TGF-β production from tissue resident macrophages and other sources inhibits immune responses, either directly by inhibiting the functions of Th1 and Th2 CD4+ effector cells and natural killer (NK) cells, or indirectly by promoting regulatory T cells (Treg) generation and function^79–81^. These examples demonstrate the opportunity for further investigation into the involvement of other immune cell types by which skeletal muscle in D2.*mdx* mice were remodeled by adiponectin receptor agonism. Overall, the diverse responses in inflammatory factors may be related to the dynamic interplay between early disease progression in the diaphragm and the rapid development and remodeling of muscle that occurs in young D2.*mdx* mice. In tibialis anterior, ALY688-SR attenuated TNF-α mRNA and protein suggesting a possible direct effect in fibres of this muscle. As there was little fibrosis and no apparent atrophy in this muscle, these results may add further credence to determining whether such early effects of treatment translate into longer-term prevention of the eventual pathology in D2.*mdx* mice that likely manifests in other muscles at later stages of disease progression. For example, atrophy of tibialis anterior fibres was reported as early as 12 weeks ^24^.

### Reduced diaphragm fibrosis and atrophy are related to attenuated mitochondrial redox stress

Mitochondrial H_2_O_2_ emission in diaphragm muscle fibres was reduced with ALY688-SR treatment. This assessment was performed with NADH-generating substrates which specifically target Complex I of the electron transport chain. The findings add to previous reports that Complex I is dysfunctional in muscle from *mdx* mouse strains ^27, 50, 82^.

The mechanism of lower Complex I-supported H_2_O_2_ emission was related to mitochondrial responsiveness to ADP. When ATP is used in muscle by ATP-hydrolyzing pathways generally located outside the mitochondria, as occurs at basal rates in resting/sedentary muscle or at increasing rates during progressively greater muscle contractions, the resulting ADP product cycles back to mitochondria to combine with inorganic phosphate to re-synthesize ATP. The resulting entry of protons from the inner membrane space through ATP synthase into the matrix lowers mitochondrial membrane potential. This event triggers a dual response in the electron transport chain whereby proton pumping from matrix to inner membrane space ensues in tandem with forward electron flux from NADH (Complex I) or other reducing equivalents at other sites. This forward electron flow reduces the probability of premature electron ‘slip’ onto oxygen to generate superoxide and its dismutated product, H_2_O_2_. Being membrane permeable, this reactive oxygen species can then be scavenged by mitochondrial antioxidants or emitted into the cytoplasm. In this way, a rise in energy demand manifested as ADP ultimately lowers H_2_O_2_ emission. The discovery reported herein reveals that *in vivo* administration of ALY688-SR enhanced the ability of ADP to attenuate mH_2_O_2_ assessed *in vitro* in both diaphragm and quadriceps of D2.*mdx* mice. However, the ability of ADP to stimulate respiration was not altered by ALY688-SR for reasons that are unclear. Nevertheless, this finding suggests that adiponectin receptor regulates some component of the phosphorylation system related to ADP’s control of bioenergetics, but the manner by which this occurs either directly or indirectly requires further investigation. The precise mechanism by which mitochondrial H_2_O_2_ contributes to atrophy and fibrosis in muscle is likely multi-factorial but is increasingly associated with myopathy in muscular dystrophies as discussed elsewhere ^13^.

Lastly, mitochondria can also export phosphate in the form of phosphocreatine with creatine cycling back for re-phosphorylation by ATP in the matrix. This pathway was assessed by including creatine in the media during assessments and compared to assessments in the absence of creatine as discussed above. As similar conclusions were made in both assay conditions, the results do not suggest that ALY688-SR rescue mitochondrial creatine metabolism in this study design^51^.

### The relationship between histopathology and muscle force responses to ALY688-SR

The attenuation of diaphragm fibrosis and atrophy occurred concurrently with lower *in vitro* force production. The reason for this lower force production response to short-term treatment of the drug is not apparent but may be related to several factors. First, treatment commenced at 7 days of age up to 28 days of age. This period of development involves rapid remodeling of muscle tissue that is regulated by extensive coordination of fibroblasts and immune cells. As these cell types are known to be influenced by adiponectin ^83, 84^, it becomes difficult to predict how adiponectin agonists may influence the balance between muscle mass, extracellular matrix components such as collagen and, force generating capacity. Second, this dynamic period of remodeling is exacerbated by the presence of early-stage disease stressors arising from a lack of dystrophin expression in this model. While speculative, when considering the increased work of breathing that occurs in people with muscular dystrophies ^85^, we did not determine whether the reduced *in vitro* force production in diaphragm was a result of reduced work of breathing following drug treatment. In line with this speculation, adiponectin-deficient mice have lower dynamic lung compliance ^86^ predicting that a greater amount of force would be required to increase lung volume. Whether this means that adiponectin lowers the work of breathing remains to be determined but this possibility could explain whether the lower diaphragm force in response to ALY688-SR was an adaptive response to lower demands. However, the reduced recovery from fatigue would not align with this and remains difficult to explain. The attenuation of diaphragm fibrosis and atrophy nonetheless align with clinical interest in these specific outcomes, and their preservation seen herein points to a need for further investigation to determine if longer-term treatment eventually preserves force production once adolescence is complete.

In summary, these results suggest that the recently developed adiponectin receptor agonist ALY688-SR prevents much of the fibrosis in diaphragm and partially prevents atrophy in multiple fibre types. These effects were observed after 3 weeks of short-term treatment in young mice at early stages of the disease and was related to dynamic changes in inflammatory markers and attenuated mitochondrial H_2_O_2_ emission – a known metabolic stressor that occurs in D2.*mdx* muscle. These results warrant further investigation into the longer-term effects of ALY688-SR on other muscles that eventually develop fibrosis and atrophy as well as the relationship of these effects to force production in limb muscle and diaphragm.

## Online supplemental material

Additional supporting information pertaining to methodologies and results are available online.

## Data Availability Statement

The data that support the findings of this study are available on request from the corresponding author.

## Author Contributions

Study design and conception were prepared by C.A. Bellissimo, S. Gandhi, G. Sweeney, C.G.R. Perry and Allysta Pharmaceuticals. Experimental work and analyses were performed by C.A. Bellissimo, S. Gandhi, L.N. Castellani, M. Murugathasan, L.J. Delfinis, A. Thuan, M.C. Garibotti, Y. Seo and I.A. Rebalka. The manuscript was written by C.A. Bellissimo. and C.G.R. Perry. All authors contributed to the manuscript and approved the final version.

## Nonstandard Abbreviations

AdipoR: adiponectin receptor
ADP: adenosine diphosphate
ATP: adenosine triphosphate
CDNB: 2,4-dinitrochlorobenzene
CSA: cross sectional area
D2.*mdx*: murine model of DMD on DBA background
DMD: Duchenne muscular dystrophy
FADH_2_: flavin adenine dinucleotide
IL: 1β-interleukin 1-beta
IL: 6-interleukin 6
*mdx*: murine model of DMD on C57/Bl10 background
MFD: minimal feret diameter
mH_2_O_2_: mitochondrial hydrogen peroxide
MHC: myosin heavy chain
NADH: nicotinamide adenine dinucleotide
NK: natura killer cells
O_2_^•-^: superoxide
TGF-β: transforming growth factor beta
TNF-α: tumor necrosis factor alpha
Treg: regulatory T cells

## Acknowledgements

ALY688-SR was provided by Allysta Pharmaceuticals. Funding was provided to C.G.R.P. by Allysta Pharmaceuticals, the National Science and Engineering Research Council (no. 2019-06687) and an Ontario Early Researcher Award (C.G.R.P., no. 2017-0351) with infrastructure supported by Canada Foundation for Innovation, the James. H. Cummings Foundation, and the Ontario Research Fund. Funding was provided by the National Science and Engineering Research Council to A.A.A.S. (no. 2017-05073) and T.J.H (2018-06324 and 2018-522456) and by the Canadian Institutes of Health Research to G.S. Scholarships were awarded to C.A.B. (NSERC PGS-PhD, MITACS Accelerate), L.N.C. (MITACS Accelerate), L.J.D. (NSERC CGS-D), S.G. and A.T. (Ontario Graduate Scholarships).

