## Supplemental Data for "The adiponectin analogue ALY688-SR attenuates diaphragm fibrosis, atrophy and mitochondrial stress in a mouse model of Duchenne muscular dystrophy"

**Supplemental Results**

**Supplemental Table 1: Anthropometrics effects of ALY688-SR treatment**

| | **Wild-Type** | **D2.*mdx*-VEH** | **D2.*mdx*-LD** | **D2.*mdx*-HD** |
|---|---|---|---|---|
| *Body Weight (g)* | 15.8 ± 2.6 | 13.2 ± 1.5* | 12.4 ± 1.3$^{\#}$ | 13.3 ± 1.6†^ |
| *Tibial Length (mm)* | 15 ± 0.7 | 14 ± 0.6* | 14 ± 0.5$^{\#}$ | 14 ± 0.5† |
| **Muscle (mg/mm)** | | | | |
| *Quadriceps* | 6.4 ± 1.0 | 6.7 ± 1.3 | 6.1 ± 0.7 | 6.7 ± 0.9 |
| *Extensor Digitorum Longus* | 0.3 ± 0.1 | 0.3 ± 0.1 | 0.3 ± 0.1 | 0.3 ± 0.1 |
| *Tibialis Anterior* | 1.7 ± 0.2 | 1.7 ± 0.3 | 1.6 ± 0.3 | 1.6 ± 0.3 |
| *Soleus* | 0.3 ± 0.1 | 0.3 ± 0.1 | 0.3 ± 0.1 | 0.3 ± 0.1 |
| *Plantaris* | 0.5 ± 0.1 | 0.5 ± 0.2 | 0.5 ± 0.1 | 0.5 ± 0.1 |
| *Gastrocnemius* | 3.8 ± 0.4 | 3.6 ± 0.5 | 3.5 ± 0.3 | 3.6 ± 0.4 |
| *Triceps* | 3.1 ± 0.8 | 3.5 ± 1.3 | 3.3 ± 0.8 | 3.8 ± 0.9† |
| **Adipose Pad (mg/mm)** | | | | |
| *Inguinal* | 7.2 ± 2.0 | 3.1 ± 1.1* | 2.9 ± 0.7$^{\#}$ | 3.6 ± 0.8† |
| *Epididymal* | 8.9 ± 3.7 | 1.5 ± 0.5* | 1.1 ± 0.4$^{\#}$ | 1.8 ± 0.6† |
| **Viscera (mg/mm)** | | | | |
| *Spleen* | 5.4 ± 1.2 | 5.0 ± 1.3 | 4.3 ± 1.07$^{\#‡}$ | 5.1 ± 1.0^ |
| *Kidney* | 8.3 ± 1.9 | 7.1 ± 1.4* | 6.6 ± 1.0$^{\#}$ | 7.3 ± 1.0† |
| *Liver* | 55.0 ± 7.5 | 55.7 ± 9.4 | 48.1 ± 4.6$^{\#‡}$ | 59.2 ± 9.0†^ |

Results represent mean ±SD; n=7-10. All p values are FDR-adjusted by Benjamini, Krieger and Yekutieli post-hoc analyses. *p<0.05 WT vs D2.*mdx*- VEH; #p<0.05 WT vs D2.*mdx*- LD; †p<0.05 WT vs D2.*mdx*- HD; ‡p<0.05 D2.*mdx*-VEH vs D2.*mdx*- LD; §p<0.05 D2.*mdx*-VEH vs D2.*mdx*-HD; ^p<0.05 D2.*mdx*-LD vs D2.*mdx*-HD

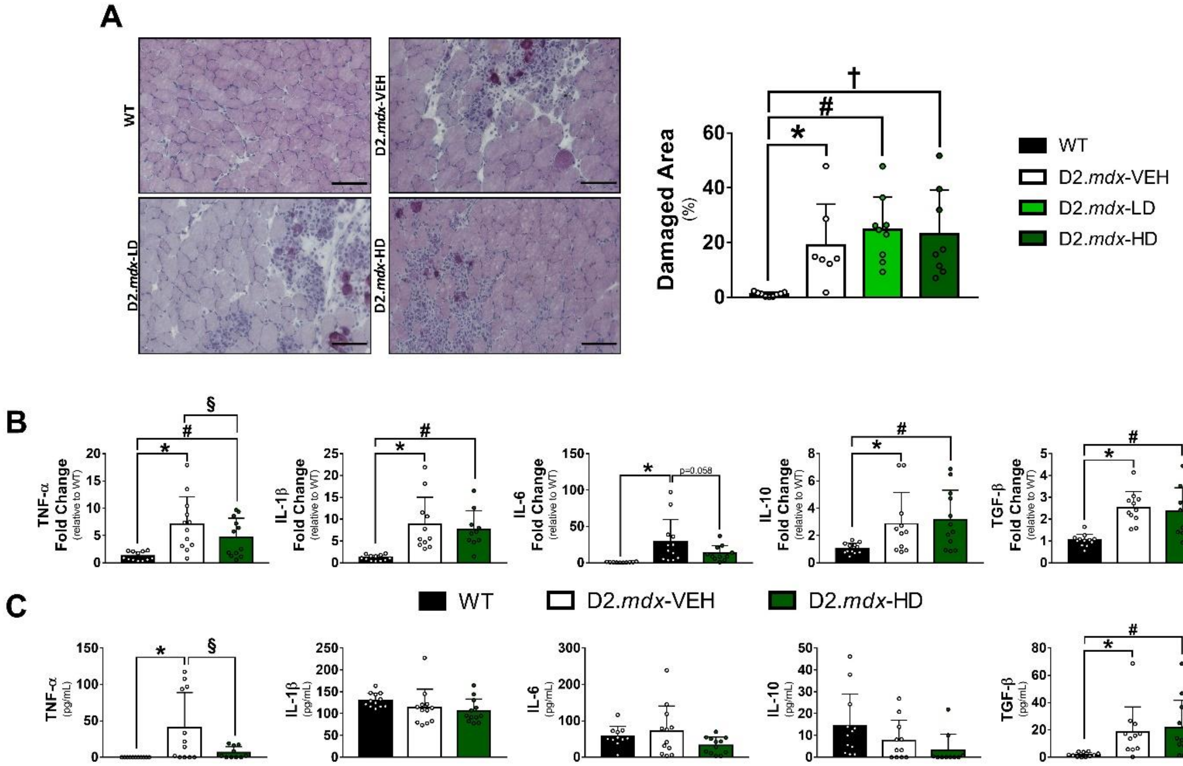

**Supplemental Figure 1: Effects of ALY688 on inflammation and muscle damage in D2.*mdx* tibialis anterior**. (A) Hematoxylin & eosin staining were used to assess areas of muscle damage which includes areas of necrosis and fibrosis and expressed as a percentage of total area. Scale bar =100µm. (B) qPCR were used to assess mRNA fold changes of TNF-α, IL-1β, IL-6, IL-10 and TGF-β across WT and D2.*mdx* mice treated with vehicle or high dose ALY688 treatment. qPCR results were normalized to *Rplp0*, and the subsequent ratios were presented as relative expression compared with WT values. (C) Protein levels of the same cytokines, were assessed by BioLegend Multiplex using flow cytometry. Results represent mean ±SD; n=7-12. All p values are FDR-adjusted by Benjamini, Krieger and Yekutieli post-hoc analyses. *p<0.05 WT vs D2.*mdx*- VEH; #p<0.05 WT vs D2.*mdx*- LD; †p<0.05 WT vs D2.*mdx*- HD; §p<0.05 D2.*mdx*-VEH vs D2.*mdx*-HD.

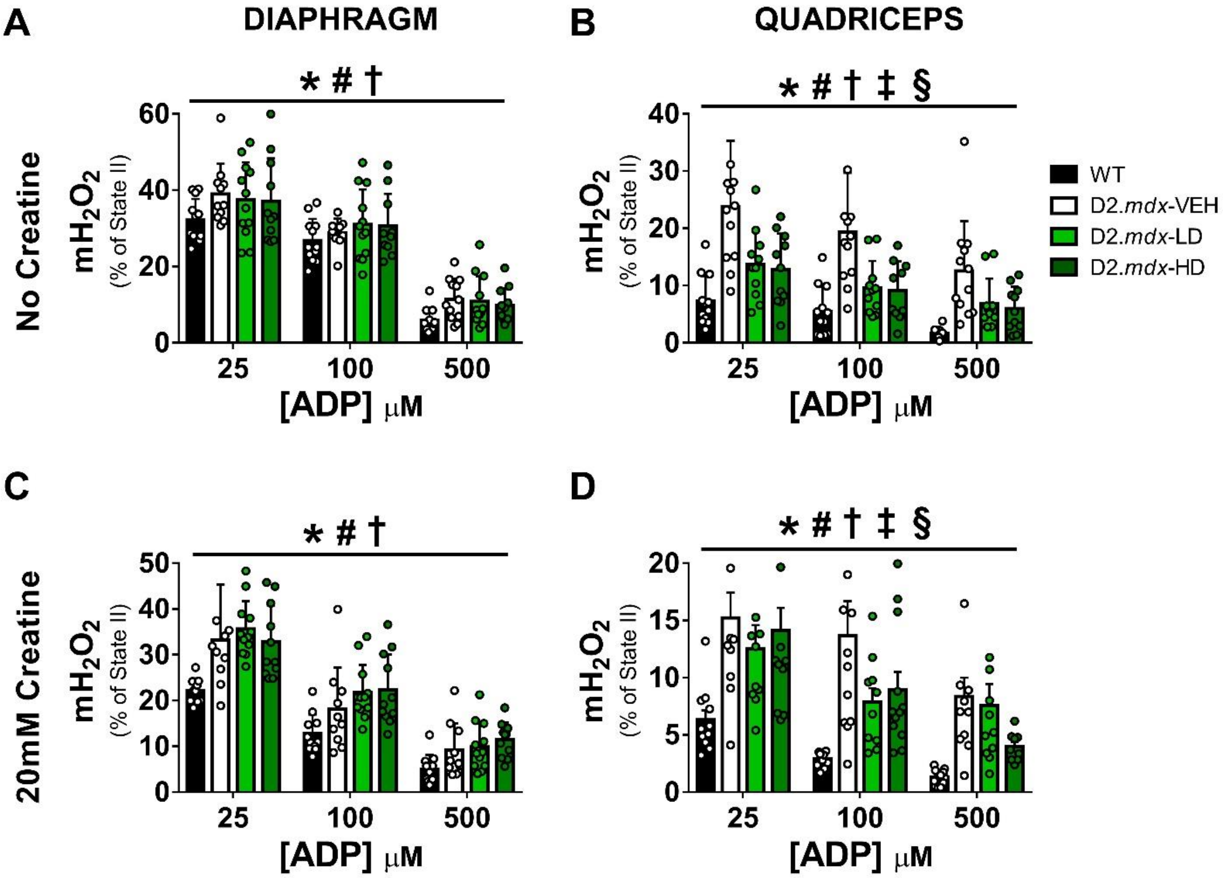

**Supplemental Figure 2**: **ALY688-SR enhances ADP attenuation of mitochondrial $H_2O_2$ emission supported by succinate in quadriceps.** The ability of ADP to attenuate mitochondrial $H_2O_2$ emission ($mH_2O_2$) stimulated by succinate was assessed in permeabilized fibres of diaphragm (A,B) and quadriceps (C,D). 10mM succinate ($FADH_2$) was used to stimulate electron flow in reverse direction from complex II to complex I to generate superoxide that is dismutated to $H_2O_2$. ADP's attenuation of $mH_2O_2$ was assessed in the absence (A,C) and presence (B,D) of creatine (accelerator of matrix ADP/ATP cycling) across a range of ADP concentrations depicting increasing metabolic demand [1-3]. Analyses were performed in the diaphragm (A,B) and quadriceps (C,D). Results represent mean ±SD; n=10-12. All p values are FDR-adjusted by Benjamini, Krieger and Yekutieli post-hoc analyses. *$p<0.05$ WT vs D2.mdx- VEH; #$p<0.05$ WT vs D2.mdx- LD; †$p<0.05$ WT vs D2.mdx- HD; ‡$p<0.05$ D2.mdx-VEH vs D2.mdx- LD; §$p<0.05$ D2.mdx-VEH vs D2.mdx-HD.

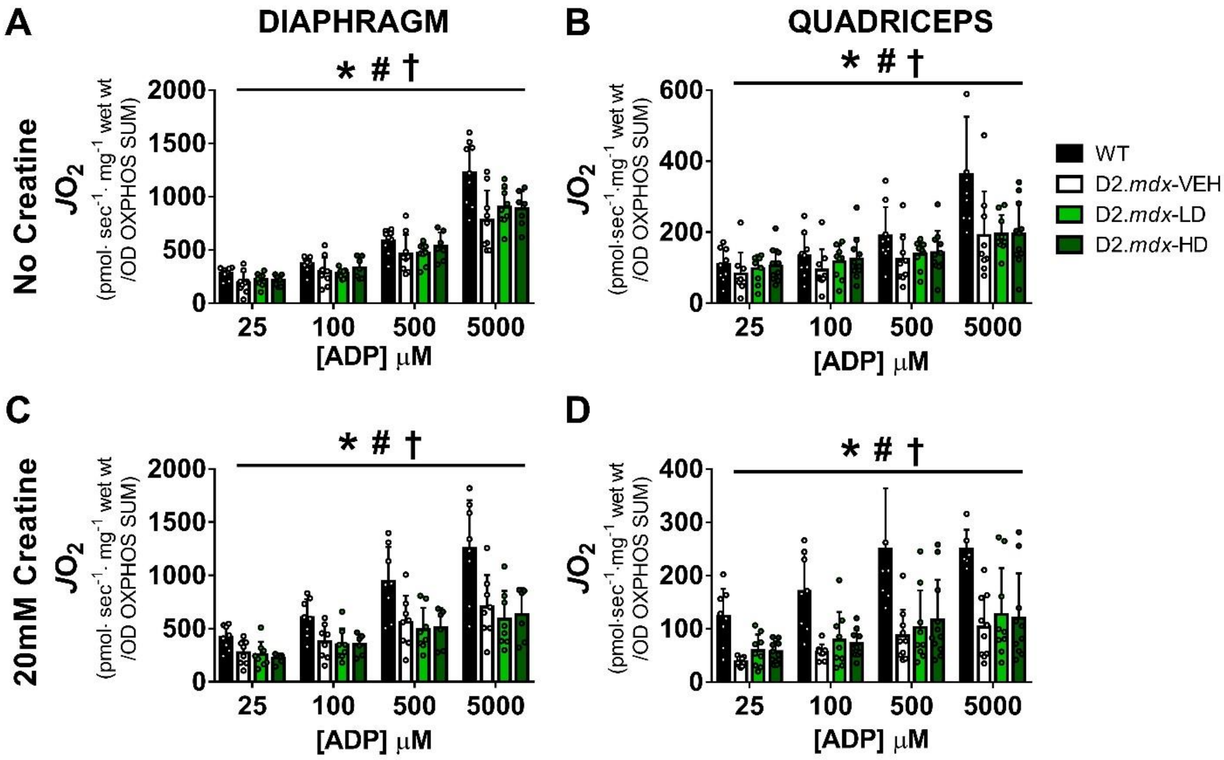

**Supplemental Figure 3: Mitochondrial respiration normalized to mitochondrial content markers is not altered by ALY688-SR treatment.** Complex-I supported respiration (assessed by oxygen flux, $JO_2$) normalized to protein markers of the electron transport chain (from Figure 6B) as an index of mitochondrial content in diaphragm (A,B) and quadriceps (C,D) in the absence (A,C) and presence (B,D) of 20mM creatine (to accelerate ADP/ATP cycling) across a range of increasing metabolic demand (ADP concentrations) [7, 15, 16]. Results represent mean ±SD; n=9-12. All p values are FDR-adjusted by Benjamini, Krieger and Yekutieli post-hoc analyses. *p<0.05 WT vs D2.*mdx*- VEH; #p<0.05 WT vs D2.*mdx*- LD; †p<0.05 WT vs D2.*mdx*- HD.

**Supplemental Table 2: Reductions in creatine-accelerated mitochondrial respiration supported by pyruvate and succinate are not altered by ALY688-SR.**

| | Diaphragm | | | | Quadriceps | | | |
|---|---|---|---|---|---|---|---|---|
| **[ADP]** (µM) | WT | D2.*mdx*-VEH | D2.*mdx*-LD | D2.*mdx*-HD | WT | D2.*mdx*-VEH | D2.*mdx*-LD | D2.*mdx*-HD |
| **Creatine-dependent Complex I-supported respiration (pyruvate)** (pmol $O_2$·sec$^{-1}$·mg$^{-1}$ wet weight) | | | | | | | | |
| **0 (State II)** | 39.9± 12.0 | 21.4± 8.2* | 19.4± 9.9# | 20.4± 4.4† | 12.5± 4.8 | 5.0± 1.8* | 5.8± 3.1# | 7.0± 2.5† |
| **25** | 55.4± 15.9 | 30.9± 13.4* | 21.6± 5.0#,‡ | 27.7± 5.5† | 25.3± 6.7 | 8.0± 3.4* | 9.8± 5.3# | 10.15± 6.0† |
| **100** | 81.0± 25.4 | 45.3± 17.7* | 30.5± 7.2#,‡ | 42.2± 9.0† | 35.8± 9.1 | 11.5± 4.5* | 13.4± 8.4# | 12.6± 7.3† |
| **500** | 126. 7± 45.4 | 68.4± 28.2* | 50.2± 18.3# | 64.1± 20.2† | 54.2± 13.3 | 15.1± 7.1* | 17.8± 10.6# | 17.3± 10.7† |
| **5000** | 174.2± 66.0 | 84.7± 34.9* | 62.0± 23.2# | 85.7± 34.8† | 74.6± 22.5 | 18.7± 10.0* | 22.2± 12.9# | 22.0± 16.2† |
| **Creatine-dependent Complex I+II-supported respiration (pyruvate, glutamate, succinate)** (pmol $O_2$·sec$^{-1}$·mg$^{-1}$ wet weight) | | | | | | | | |
| **Max** | 215.3± 79.7 | 161.7± 55.1# | 135.3± 37.7 | 159.9± 33.8 | 122.7± 51.7 | 60.3± 17. 6* | 60.1± 21.0# | 64.9± 19.1† |

Creatine accelerates matrix-ADP/ATP cycling through mitochondrial creatine kinase activation [3-9] in contrast to data in Figure 6 which was assessed in the absence of creatine (slower state of ADP/ATP cycling). Complex-I supported respiration stimulated by NADH generation (5mM pyruvate and 2mM malate) in permeabilized fibres across a range of ADP concentrations representing increasing states of metabolic demand: 25µM (modeling resting muscle [2]); 100µM (modeling high intensity exercise [1,6]), 500uM (maximal) and 5000µM (supramaximal) was assessed in the diaphragm (A) and quadriceps (B). Following ADP titrations, 20mM succinate was added to generate $FADH_2$ which stimulates forward electron flow (due to presence of ADP) from Complex II (yielding Complex I+II supported respiration). Results represent mean ±SD; n=9-12. All p values are FDR-adjusted by Benjamini, Krieger and Yekutieli post-hoc analyses. *p<0.05 WT vs D2.*mdx*- VEH; #p<0.05 WT vs D2.*mdx*- LD; †p<0.05 WT vs D2.*mdx*- HD; ‡p<0.05 D2.mdx-VEH vs D2.mdx- LD.

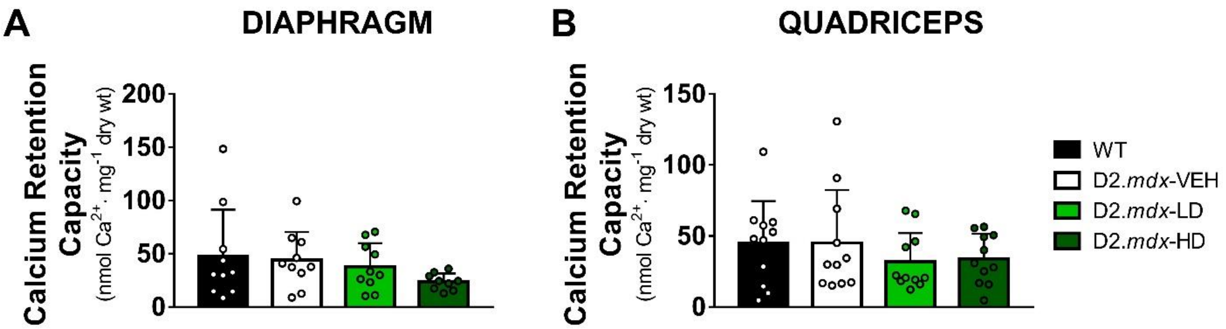

**Supplemental Figure 4: Calcium-induced mitochondrial permeability transition pore opening in D2.*mdx* mice after ALY688-SR.** Calcium retention capacity was measured spectrofluorometrically by titrating calcium chloride into permeabilized muscle fibres from diaphragm (A) and quadriceps (B) until mitochondrial permeability transition pore opening was triggered. Results represent mean ±SD; n=9-11.

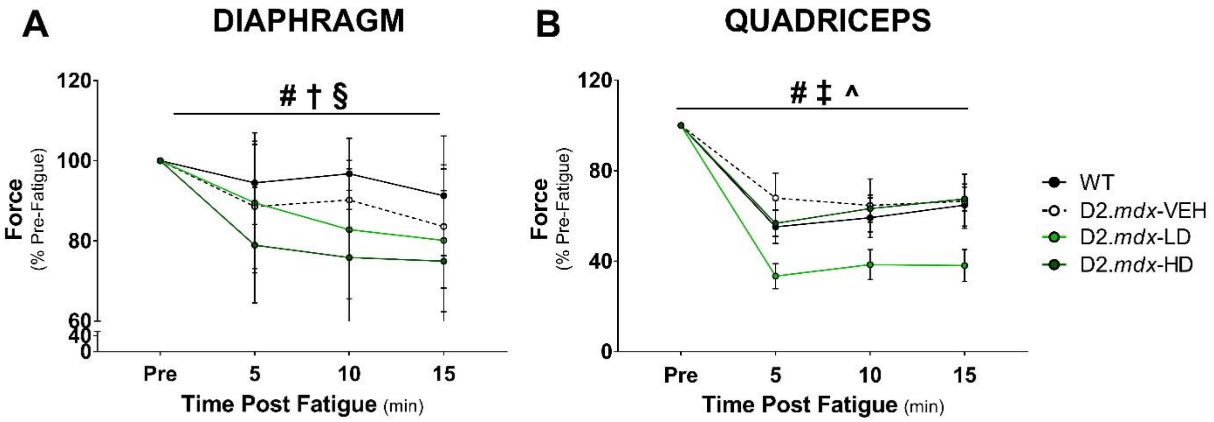

**Supplemental Figure 5: Effects of ALY688-SR treatment on recovery after muscle fatigue in D2.*mdx* mice**. *In-vitro* diaphragm strips (A) and *in-situ* quadriceps (B) underwent fatiguing protocols (70 Hz for 350 ms every 2 seconds for 5minutes). Prior to fatigue, 5-,10- and 15 minutes post fatigue, single stimulations were used to assess recovery and expressed as a % of pre-fatigue contraction. Results represent mean ±SD; n=8-11. All p values are FDR-adjusted by Benjamini, Krieger and Yekutieli post-hoc analyses. *p<0.05 WT vs D2.*mdx*- VEH; #p<0.05 WT vs D2.*mdx*- LD; †p<0.05 WT vs D2.*mdx*- HD; ‡p<0.05 D2.*mdx*-VEH vs D2.*mdx*- LD; §p<0.05 D2.*mdx*-VEH vs D2.*mdx*-HD; ^p<0.05 D2.*mdx*-LD vs D2.*mdx*-HD
